# Persistence of the ecological niche in pond damselflies underlies a stable adaptive zone despite varying selection

**DOI:** 10.1101/2022.10.19.512907

**Authors:** Anjali Gupta, Erik I. Svensson, Henrik Frietsch, Masahito Tsuboi

## Abstract

Following the development of regression-based methods to estimate natural and sexual selection, evolutionary biologists have quantified the strength, mode and direction of selection in natural populations. Although this approach has been successful, its limitations include lack of replication across species, compromising the generality of the inferences beyond microevolutionary time scales. Here, we carried out a comparative study of selection on wing shape and body size across multiple populations of two closely related and ecologically similar pond damselflies: *Enallagma cyathigerum* and *Ischnura elegans* (Odonata: Coenagrionidae). We found weak stabilizing selection on wing shape in both sexes, and no evidence that selection on this trait differed between the species. In contrast, selection on body size was curvilinear in males and directional in females, and they differed in form (males) and intensity (females) between these two species. By analyzing selection on the fine-grained spatial scale, we found that selection on male body size was shaped by the local mating system, and the relationship between mating system characteristics and directional selection was remarkably consistent across these species. Finally, we present a graphical model that links contemporary selection and macroevolution. Based on this model, we conclude that the persistence in ecological modes of life in pond damselflies offers a plausible explanation for why varying selection in nature may still result in a stable adaptive zone lasting millions of years.

## Introduction

Since Russell Lande and Stevan Arnold introduced regression-based statistical methods to quantify natural and sexual selection (Lande & Arnold, 1983), evolutionary biologists have sought to describe and understand the causes of phenotypic selection in nature (Endler, 1986, Mitchell-Olds & Shaw, 1987, Wade & Kalisz, 1990, Kingsolver et al., 2001). This has resulted in a rich scientific literature about the strength, mode and direction of selection in wild populations of animals and plants (Siepielski et al., 2017, Kingsolver et al., 2001, Kingsolver & Diamond, 2011, Siepielski et al., 2009). However, several methodological and conceptual problems remain (Mitchell-Olds & Shaw, 1987, Wade & Kalisz, 1990, Hereford et al., 2004, De Lisle & Svensson, 2017, Svensson, 2023), including the lack of our understanding of whether and how estimates of phenotypic selection measured at contemporary populations are related to divergence across populations, species, or higher taxa (e.g., macroevolution). The adaptive landscape has been proposed as a conceptual framework that unifies phenotypic selection within populations (i.e., microevolution) with macroevolution (Arnold et al., 2001, Svensson & Calsbeek, 2012, Arnold, 2023). Under this framework, populations are conceptualized as evolving on fitness surfaces, where they are subject to selection, genetic drift and mutation (Lande, 1976). Populations are expected to climb the closest adaptive peak in the direction where short-term fitness is maximized (Wright, 1932) whereas genetic correlations can bias the evolutionary trajectories towards directions with high amount of additive genetic variance (Schluter, 1996, Walsh & Blows, 2009). Estimates of selection coefficients capture local landscape topography, which can be used to infer the selection surface occupied by panmictic populations (Fear & Price, 1998, Gavrilets, 2004, Schluter, 2000) and the dynamics of the landscape itself (de Villemereuil et al., 2020, Chevin et al., 2015). There are ongoing attempts to synthesize this line of largely conceptual research with empirical data from wild populations (Beausoleil et al., 2023, Stroud et al., 2023).

In a seminal paper, Arnold et al. (2001) envisioned that the integration of microevolution and macroevolution requires a shift from a static to a dynamic view of adaptive landscape. It triggered an important conceptual shift in the literature of phylogenetic comparative studies (Estes & Arnold, 2007, Uyeda et al., 2011, Hansen et al., 2008) where the parameters of an Ornstein-Uhlenbeck model began to be recognized as descriptors of the dynamics of adaptive zone—a collection of adaptive landscapes inhabited by populations and species that share an ecological mode of life (Simpson, 1944, Hansen, 1997)—rather than the dynamics of species evolving on a fixed adaptive landscape. Thus, the adaptive landscape estimated from present-day populations and the adaptive zone estimated from a dataset that contains multiple species can be conceptually related to study the link between contemporary selection and macroevolution. However, this idea is currently considered a metaphor with limited bearings on empirical research, because many previous studies have shown that phenotypic selection often fluctuates over short time scales (Kinnison & Hendry, 2001, Gosden & Svensson, 2008, Grant & Grant, 2002, Siepielski et al., 2017) and stabilizing selection is not as ubiquitous in nature as have been assumed in many theoretical models (Kingsolver & Diamond, 2011, Hereford et al., 2004). Such findings are at odds with the core premise of the adaptive zones—the assumption that adaptive phenotypes are stable over millions of years. Therefore, to construct an operational research program of the adaptive landscape “*as a conceptual bridge between micro- and macroevolution*” (Arnold et al., 2001), we need to elucidate how perpetually moving adaptive landscapes at the population level may yield a stable adaptive zone lasting millions of years.

The persistence of environmental and ecological conditions (i.e., ecological niche) has long been a major hypothesis to explain the stability of adaptive zone. In the classic work on the evolution of high-crown cheek teeth (hypsodonty) in horses, Simpson (1944) argued that the evolution of hypsodonty in the early to middle Miocene reflects the spread of Savanna and grassland habitats during this era. A microevolutionary analogue to this argument is abound in the literature of phenotypic selection. For example, it has been suggested that trait-fitness covariances (i.e., selection gradients) are not necessarily causal and should be viewed as preliminary hypotheses in need of experimental verification through trait manipulations (Mitchell-Olds & Shaw, 1987). Furthermore, it has also been argued that functional analysis of selective agents or detailed field observations of wild populations are necessary to identify environmental agents, demographic factors and ecological causes behind observed selection patterns (Svensson & Sinervo, 2000, MacColl, 2011, Opedal, 2021). These body of work suggest that ecology can provide critical insights to bridge micro- and macroevolution. Ecological causes behind phenotypic selection in natural populations has been identified in a handful of cases (Arnqvist, 1992, Grant & Grant, 2002, Gosden & Svensson, 2008, Martin, 2016), but its macroevolutionary relevance has rarely been explored (Punzalan & Rowe, 2016).

The insect order Odonata (damselflies and dragonflies) is a popular system for studies of phenotypic selection because a rich background knowledge of natural history (Corbet, 1999) enables characterization of the adaptive landscape in different environmental and ecological contexts. Here, we study two ecologically and phenotypically similar sympatric damselfly species (Fig. S1): the common bluetail damselfly (*Ischnura elegans*) and the common bluet (*Enallagma cyathigerum*). These two species and other members of the family Coenagrionidae (pond damselflies) are ecologically similar, show evidence of neutral community dynamics and may therefore fluctuate in abundances due to ecological drift (McPeek & Brown, 2000, Siepielski et al., 2010). Molecular phylogenetic evidence suggest that these two genera have been separated by >12 million years (Willink et al., 2024b, Blow et al., 2021). The high ecological similarity of these two closely related species from two different genera provide a unique opportunity to investigate if the adaptive zone has remained stable for millions of years, while simultaneously considering variation in the local adaptive landscapes that vary on fine-grained spatial and temporal scales. Previous selection studies suggest that spatial variation in phenotypic selection in *I. elegans* is partly driven by the dynamics of color polymorphism in females (Gosden & Svensson, 2008). These female color morphs experience negative frequency-dependent selection that maintains the polymorphism on micro- and macroevolutionary time scales (Le Rouzic et al., 2015, Svensson et al., 2005, Willink et al., 2024a). We hypothesize that spatially and temporally varying short-term selection in these damselflies may result in a stable adaptive zone over macroevolutionary time scales because their shared ecological mode of life imposes similar selective pressures on both species.

We have three objectives in this study. First is to estimate univariate and multivariate selection on two important fitness-related traits—body size and wing shape—and compare these within and between species. Second is to quantify demographic and mating system parameters across multiple populations of these two species. These demographic and mating system parameters include the opportunity for sexual selection (Arnold & Wade, 1984), mating frequencies, male and female densities, operational sex ratio (OSR) and the frequency of male-mimicking female color morphs (“androchrome females”). These mating system parameters provide information about the local sexual selection regimes, male-male competition (opportunity for sexual selection, OSR, male density) and the intensity of male mating harassment and sexual conflict (mating frequencies and the frequency of androchrome female morphs; see Blow et al., 2021, Takahashi et al., 2014, Gosden & Svensson, 2009, Svensson & Abbott, 2005). Finally, by combining information from these different estimates about the ecological causes of selection, we aim to understand and explain how selection that varies in contemporary time scales may result in a stable adaptive zone in two ecologically similar pond damselflies.

## Materials & Methods

### Study system

We studied two species of damselflies, the common bluet, *Enallagma cyathigerum* and the common bluetail damselfly, *Ischnura elegans* in the family Coenagrionidae (Odonata; Zygoptera). These species are generalist predators that feed on small flying insects and they occur in diverse aquatic habitats, typically open landscapes in ponds, lake shores, and slow flowing streams (Smallshire & Swash, 2020). Females of both species are polymorphic; females of *E. cyathigerum* exhibit two color morphs: androchrome (mature females with blue thorax mimicking males; Fig. S1A *ii*) and gynochrome (mature females with olive-green thorax; Fig. S1A *iii*), while females of *I. elegans* exhibit three heritable color morphs during the adult stage: androchrome (mature females with blue thorax; Fig. S1A *v*), *infuscans* (Fig. S1A *vi*) and *infuscans-obsoleta* (Fig. S1A *vii*). Mating system of both species are characterized by scramble male-male competition, where non-territorial males rest or fly near water bodies, chase females that water for oviposition and compete for the opportunity for mating (Corbet, 1999). A male that successfully clasps a female will form the “tandem position” (Fig. S1E), followed by the formation of “a mating wheel” (Fig. S1D) during which egg fertilization takes place (Corbet, 1999).

### Field work

Field surveys were carried out to collect *I. elegans* and *E. cyathigerum* at 20 different field sites in and around Lund, Sweden. Field sampling was conducted at small ponds in the following localities: Borgeby, Bunkeflostrand, Flackarp, Flyinge 30 A1, Flyinge 30 A3, Genarp, Gunnesbo, Hoje A14, Hoje A6, Hoje A7, Habo Gard, IKEA, Ilstorp, Krutladan, Ladugårdsmarken, Lomma, Lunnarp, Råbydammen, Vombs vattenverk, and Vombs Bruksgård (Fig. S2). Individuals were captured using hand-held nets while slowly walking around waterbodies. Upon capture of individuals flying without a partner, we examined sex and kept males and females separately in small net cages. Individuals found as either a tandem or as a copulating couple were kept in plastic cups. We visited these populations between the hour of 08.00 and 13.00 in all partially or fully sunny days with temperature >15°C in May, June and July of 2020 and 2021. At each visit, we sampled between 20 and 30 minutes, and between 3-5 people participated in the sampling. The captured damselflies were kept in cooling bags to protect them from overheating and were brought back to the lab for recording phenotypic data and setting up mated females for egg-laying for fecundity measurements. For each individual, we recorded sex, female morph, and morphometric measurements (see subsequent section for more detail). The dataset obtained from this field survey include 58 sampling visits to capture *I. elegans* (in 2020 and 2021) and 27 sampling visits to capture *E. cyathigerum* (in 2021), and constitutes in total of 497 single males, 236 single females, and 524 mating couples of *I. elegans* and 420 single males, 181 single females, and 411 mating couples of *E. cyathigerum*. This dataset was used to estimate selection gradients (details are presented in subsequent section).

In a complementary field study, we quantified mating systems of *I. elegans* and *E. cyathigerum*. Here, we captured individuals of damselflies and dragonflies regardless of the species identity along predefined transects, with the aim of quantifying species composition, operational sex-ratio and density of local Odonata fauna. At each visit, 3-5 people sampled between 10-20 minutes, who identified and recorded all captured individuals in terms of species, sex, female color morph, sexual maturity and mating status (either captured as single or copula/tandem). These visits were carried out independently from the field work presented above. After recording this information, animals were released. From the data, we extracted data of *I. elegans* and *E. cyathigerum* from four seasons (2018-2021), that were collected during a total of 366 visits. These data constitute a total observation of 3527 single males, 1788 single females, and 630 mating couples of *I. elegans* and 3031 single males, 403 single females, and 282 mating couples of *E. cyathigerum*. This dataset was used to compare and quantify differences in mating system between these two species.

### Fitness components

We measured two fitness components: (1) mating success that characterizes sexual selection in males and (2) fecundity that characterizes natural selection in females. Sexual selection on males was estimated by comparing the phenotypes of mated and unmated males upon capture in the field. Males that were captured in tandem or in a copula were classified as ‘mated’ and assigned a mating success of ‘1’, while males that were captured solitary were classified as ‘single’ and their mating success were assigned to ‘0’. This is an established technique of quantifying mating success in damselflies (Gosden & Svensson, 2008, Steele et al., 2011). To measure female fecundity, we placed females captured in mated couples in small plastic cups with moist coffee filter to let the female lay eggs for 48hrs. The number of eggs laid by each female was subsequently counted. The number of eggs laid during this time interval likely reflects the recent ecological conditions a given female has experienced preceding oviposition, and should capture her past history of food intake rates and ambient temperatures (Svensson & Abbott, 2005). This measure of fecundity thus provides a measure of natural selection in females. Although our cross-sectional measure of fitness is often considered noisy compared to longitudinal measures such as lifetime reproductive success, this limitation is less severe in short-lived insects (Nishida, 1989) such as pond damselflies which typically live for only a few days as reproducing adults (Corbet, 1999).

### Morphometric measurements and wing image acquisition

We obtained digital images of all captured individuals using a scanner (CanoScan 5600F) at a resolution of 600 dpi to measure body size. Five linear measurements were taken for each individual to capture different aspects of the body size: total body length, thorax width, abdomen length, wing length, and width of the S4 segment (Fig. S1B). Individuals were measured from these photographs using the computer program, Fiji (Schindelin et al., 2012). All measurements were originally recorded in units of pixel, which was then converted to millimeter (mm) based on a conversion between mm and pixel. After the scan has been completed, individuals were sacrificed by exposing them to cold temperature. From sacrificed individuals, we dissected fore- and hindwings from both left and right side of the body. Dissected wings were then placed in a wet coffee filter, covered with a transparent plastic sheet, then scanned at a resolution of 2000 dpi. We obtained images for a total of 5764 wings belonging to *E. cyathigerum*, and a total of 7704 wings belonging to *I. elegans*.

### Automated assay of wing shape using ML-morph

We used a machine learning tool “ML-morph” (Porto & Voje, 2020) to measure the x-y coordinates of 17 landmark that characterize wing venation patterns (Fig. S1C). First, a training set was constructed to train ML-morph to identify the landmark positions in the wing image. We used a training set containing 400 wing images that was landmarked manually using imglab version 1.18 (compiled from Python module dlib-19.23). These 400 images subset in the training set contained a random mixture of 100 images of the 4 wings (right forewing, right hindwing, left forewing, left hindwing) of a male *I. elegans*, 100 images of a female *I. elegans*, 100 images of a male *E. cyathigerum*, and 100 images of a female *E. cyathigerum*. This training set represents variation in wing shape with respect to sex, species, and fore- or hind-wing assignment, that are considered as main source of variation in our sample set.

We used *shape.predictor* function of the ML-Morph to train the algorithm for landmarking the wing. Then, we tested the ability of ML-Morph to landmark wings accurately using *shape.tester* function, where we compared automated landmarking by ML-morph to manual landmarks, which responded to 99.2% precision in the landmarking using automation in a test set build from the training set. Finally, we applied *shape.predictor* function to landmark the remaining image set that have not been landmarked. All landmarked images were later checked manually for any errors, and inconsistent landmarks across all images were corrected. We removed 357 images (178 for *I. elegans* and 179 for *E. cyathigerum*) from the dataset before analysis because they contained an injured/broken wing where position of 1 or more landmarks could not be deduced accurately. Examination of erroneous images were done by one observer (AG).

### Geometric morphometrics and dimension reduction

The wing data obtained from ml-Morph were standardized through geometric morphometrics analysis. We performed a Generalized Procrustes Analysis (GPA) on the 17 landmarks including all measured wings using geomorph package version 4.0.5 (Adams & Otarola-Castillo, 2013). GPA transforms the landmarks by rotating, aligning, and scaling so that the resulting landmarks (i.e., aligned coordinates) describe the wing shape alone. The aligned coordinates, however, still have 30 dimensions, and dimensional reduction was necessary for further analyses. We employed two complementary approaches for dimension reduction. First, we performed a principal component analysis (PCA) on the covariance matrix containing the aligned coordinates of both species to reduce dimensionality of the major axis of morphological variation. Second, we performed a linear discriminant function analysis (LDA) on the aligned coordinates with species as a classifier using MASS package version 7.3-53. PCA and LDA were performed using the entire dataset including both species, both sexes, and both fore- and hindwings to define wing traits in a common shape space. We also performed a principal components analysis (PCA) on the covariance matrix of natural logarithmic values of the five size traits (total body length, thorax width, abdomen length, wing length, and width of the S4 segment) for both species and sexes. We log_e_-transformed the values for the size traits before running the PCA, so that the resultant PC values are mean-standardized. We averaged values obtained from left and right side of the wing of an individual whenever both sides were measured. When only one side of the wing was available, we used that available side as a representative measure. Since we found no directional asymmetry in wing shape in two species we examined (results not shown), this will unlikely create systematic bias in our measurements. In 62 individuals, we obtained scan images twice at two independent scanning sessions to evaluate measurement error due to variation in our imaging process and found that the measurement errors in aligned coordinates associated with the difference in images are on average 0.19% of the centroid size (range: 0% - 3.34%, Fig. S3). Since this level of error is negligeable compared to the effect sizes of all analyses presented in this research, it will not be accounted for in subsequent statistical analyses of wing shape.

### Quantifying mating system variation

For quantitative estimation of mating systems of these two damselfly species, we measured the following parameters: male density (total number of males caught per sampling event / total catching time, unit = number of individuals captured per minute), female density (total number of males caught per sampling event / total catching time, unit = number of individuals captured per minute), operational sex ratio (OSR, ratio of the number of mature males to the number of mature females of a species caught at a locale in a given year, unit = %), androchrome frequency (ratio of the number of the androchrome female morph of a species to the total number of females of a species whose morph was identified in one locale in one year, unit = %), mating frequency (number of males in copula / total number of males caught during one sampling event, unit = %) and the opportunity for sexual selection (*I_s_*, variance in male mating success / (average male mating success)^2^, unitless elasticity) (Arnold & Wade, 1984). We estimated *I_s_* for every locale from where we sampled populations of *E. cyathigerum* and *I. elegans*. Male mating success was defined as a binomial variable that could be either 0 or 1. We calculated variance in male mating success based on the data on the proportion of single and copulating males at each site and mean male mating success. The latter (mean male mating success) was calculated by the mean proportion of male (belonging to one species) mating at a particular locale in one field season (e.g., 0.10 if 10% are found in copula). The variance in mating success was evaluated as *p*×(1-*p*) where *p* is the probability of being found as a couple.

### Selection gradients

We used standard multiple-regression analyses to estimate the selection gradients (Lande & Arnold, 1983). Selection gradients were evaluated for body size (size-PC1) and wing shape (LD1) using mating success in the field as a male fitness component and the number of eggs laid as the female fitness component. Both linear (β) and quadratic (γ) selection coefficients were estimated for both species and sexes for all the traits using linear mixed effect models with sampling location as the random factor. The partial regression coefficients of models that includes only linear (i.e. unsquared) term were used to estimate β, while γ was estimated as the partial regression coefficients of models that includes both unsquared and squared terms. The quadratic regression coefficients and associated standard errors were multiplied by two (Stinchcombe *et al*., 2008). Note that these operations are not necessary for quadratic terms between different traits (i.e. correlational selection). The quadratic selection surfaces were visualized using cubic splines (Schluter, 1988). Fitness components (mating success for male and the number of eggs laid for females) were standardized within population and sex, by dividing individual fitness with population- and sex-specific mean fitness estimates (De Lisle & Svensson, 2017). With local fitness relativization, estimates of the global selection surface (i.e., β and γ) of a species including multiple populations approximates the adaptive landscape (Beausoleil et al., 2023, Fear & Price, 1998). Since both body size PC1 and wing shape LD1 are scaled to its own value either by taking a natural logarithm of size measurements or by performing the Generalized Procrustes Analyses, selection gradients evaluated in these traits are on the mean-standardized scale without additional transformations (see Voje et al., 2023, Lynch, 1990 for rationales behind this logic). For comparison, we also estimated the variance-standardized selection gradients by dividing them with standard deviation of the trait within species (Lande & Arnold, 1983). We also performed a bivariate model to evaluate correlational selection between LD1 and body size PC1. We analyzed forewings and hindwings separately in these analyses because LD1 of fore- and hindwings are highly correlated (r^2^ of the relationship between LD1 of fore- and hindwings is 0.95 in males and 0.94 in females).

Estimation of selection gradients and comparisons of adaptive landscape between species were made using random mixed effect models implemented in lme4 package (Bates et al., 2015). We constructed three sets of models, which all included sampling location as the random effect. First, for each sex and species separately, we modeled fitness component as the response variable and one of examined traits (size-PC1, forewing shape LD1, hindwing shape LD1) and its squared terms as the fixed explanatory variables. Second, we constructed two bivariate models, for each sex and species separately, with the same model specifications as above but include either body size PC1 and forewing shape LD1 or body size PC1 and hindwing shape LD1 and their interaction and squared terms as the fixed explanatory variables. Finally, we constructed two models for each sex separately, with the fitness component as the response variable, body size PC1 and forewing shape LD1 or body size PC1 and hindwing shape LD1 with their squared terms, species (either *I. elegans* or *E. cyathigerum*) and interaction terms between species and all traits as the fixed explanatory variables. In the last model, we identified the best model with a backward model selection procedure based on AICc (sample size-corrected Akaike Information Criterion) values.

### Selection gradients, local sexual selection regimes and mating system parameters

We evaluated the relationship between directional/quadratic selection gradients and local sexual selection regime across populations using a linear regression. For this analysis, we retained 14 populations with unique year-sampling location identity in *I. elegans* and 3 populations in *E. cyathigerum* that are represented by 20 or more samples and estimated linear (β) and quadratic (γ) selection gradients at each population as described above, but without sampling location as a random effect. Note that, throughout this study, fitness was standardized at the population level, so that the fitness surface of each location estimated here is a local representation of the adaptive landscape of corresponding species estimated in the previous section. Using estimates of β and γ from these analyses, we constructed models where trait- and sex-specific selection gradients are the response variable and one of the six variables that describe mating system (male density, female density, operational sex ratio, androchrome frequency, mating frequency, opportunity for selection) are the predictor variable and species as the fixed effect that allows species to have different intercepts. Interaction terms between predictor variables were assessed but are not presented because none were statistically significant (results not shown). In all models, inverse of standard errors of selection gradients were used to weigh observations to account for uncertainties in selection gradients. All analyses were performed in R version 4.2.2 (R Core Team, 2019) and visualization of data and results were performed using ggplot2 package version 4.0.5 (Wickham, 2016).

## Results

### Dimension reduction of body size and exploration of wing morphospace

A scatterplot of the first two principal components of five body size measurements (Fig. S4) revealed that the first PC axis explained the vast majority (93.0%) of the total variation in log of body size measurements. Based on this, we hereafter use this axis (size-PC1) as our measure of body size. A scatterplot of the first two principal components of the entire wing shape data (including both species, sex, and both fore- and hindwings) is shown in Figure S5. The first principal component (shape-PC1) explained 41.9% of the total variation, and this axis separates forewings from hindwings in terms of the width of the wing. The forewings of both species have lower PC1 values along PC1 axis, corresponding to narrower wings, while the hindwings of both species have higher values of PC1, translating to broader wings. The second principal component (shape-PC2) explained 19.2% of the total variation in wing shape, and this axis separates the two species. *E. cyathigerum* has broader proximal edge and narrower distal edge of wings as compared to *I. elegans*. Subsequent PC axes up to PC5 explained 8.9%, 6.3%, and 5.9% of total variation, respectively.

A linear discriminant function analysis (LDA) with species as a grouping factor revealed that the axis that most effectively separates wings of the two species (wing shape LD1) explains 33.3% of the total variation in wing shape and represents the variation in the width of the wing, that occurs together with the stretch of two landmarks (LM9 and LM16; Fig. S1) at the center of wing (Fig. 1). LD1 is correlated with shape-PC1 (r^2^; forewing: 62.5%, hindwing: 68.6%) and shape-PC2 (r^2^; forewing: 86.9%, hindwing: 89.0%), but not with shape PC3, PC4, and PC5 (r^2^ < 5%, Fig. S6). Within two subsets of data that include either forewings or hindwings, LD1 explained 47.1% (forewing) and 57.4% (hindwing) of the phenotypic variation within each subset, meaning that LD1 is the primary axis of phenotypic variation in these traits. To make our paper tractable, the rest of our analyses will primarily use LD1 as the representative wing shape trait. We will return to PC axes in the discussion to verify that our results and interpretations based on LD1 is applicable to other shape dimensions.

**Figure 1:**
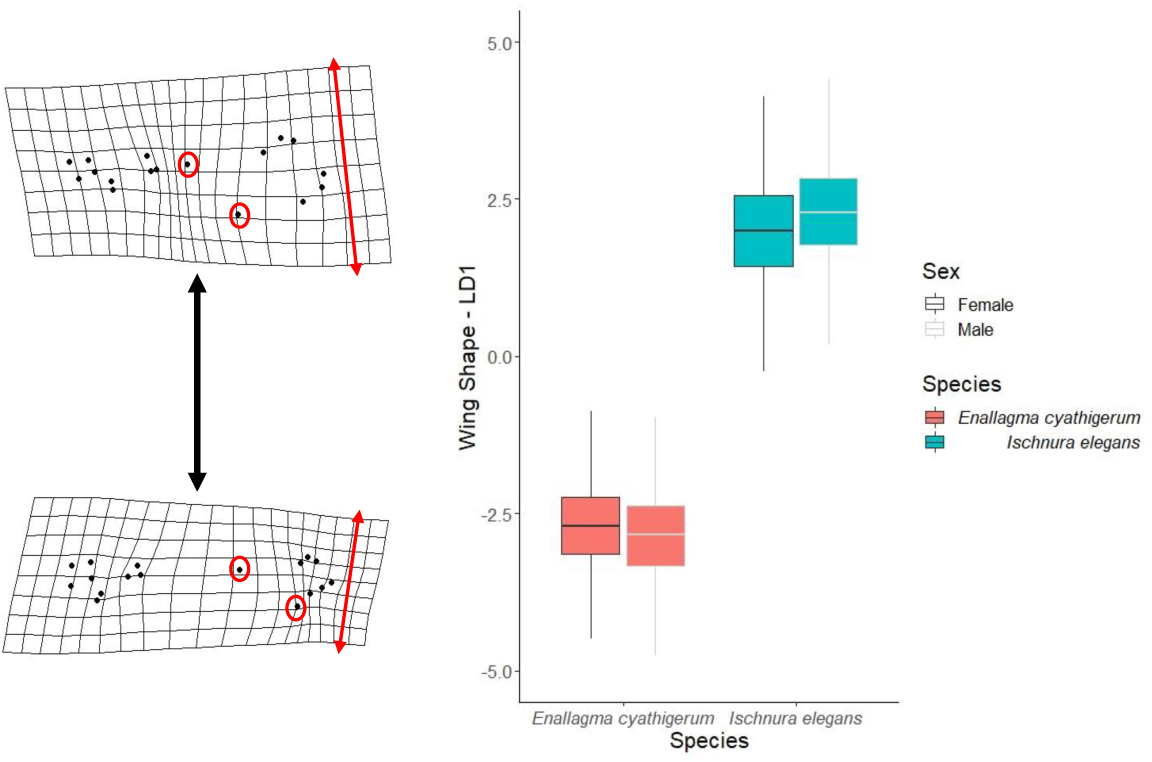
Boxplot comparing wing shape (LD1) between *I. elegans* and *E. cyathigerum.* The landmark specimens visualized in this figure represent shape associated with 2 times the minimum and maximum values of LD1.

### Selection gradients

Selection gradients evaluated at the species level are presented in Table 1 and visualized in Fig. 2. Across both species, we found that selection favors females with large body sizes. Estimates of mean-standardized directional selection gradients (β) in females (*I. elegans*: β ± SE = 0.172 ± 0.251, *E. cyathigerum*: 1.103 ± 0.700) means that doubling of body size would increase relative fitness by 17.2% in *I. elegans* and 110.3% in *E. cyathigerum*. In males, directional selection on size PC1 was weaker, whereas the quadratic selection gradients (γ) in *E. cyathigerum* indicate disruptive selection (γ ± SE = 9.226 ± 4.639) favoring either small or large individuals (Fig. 2A).

**Figure 2:**
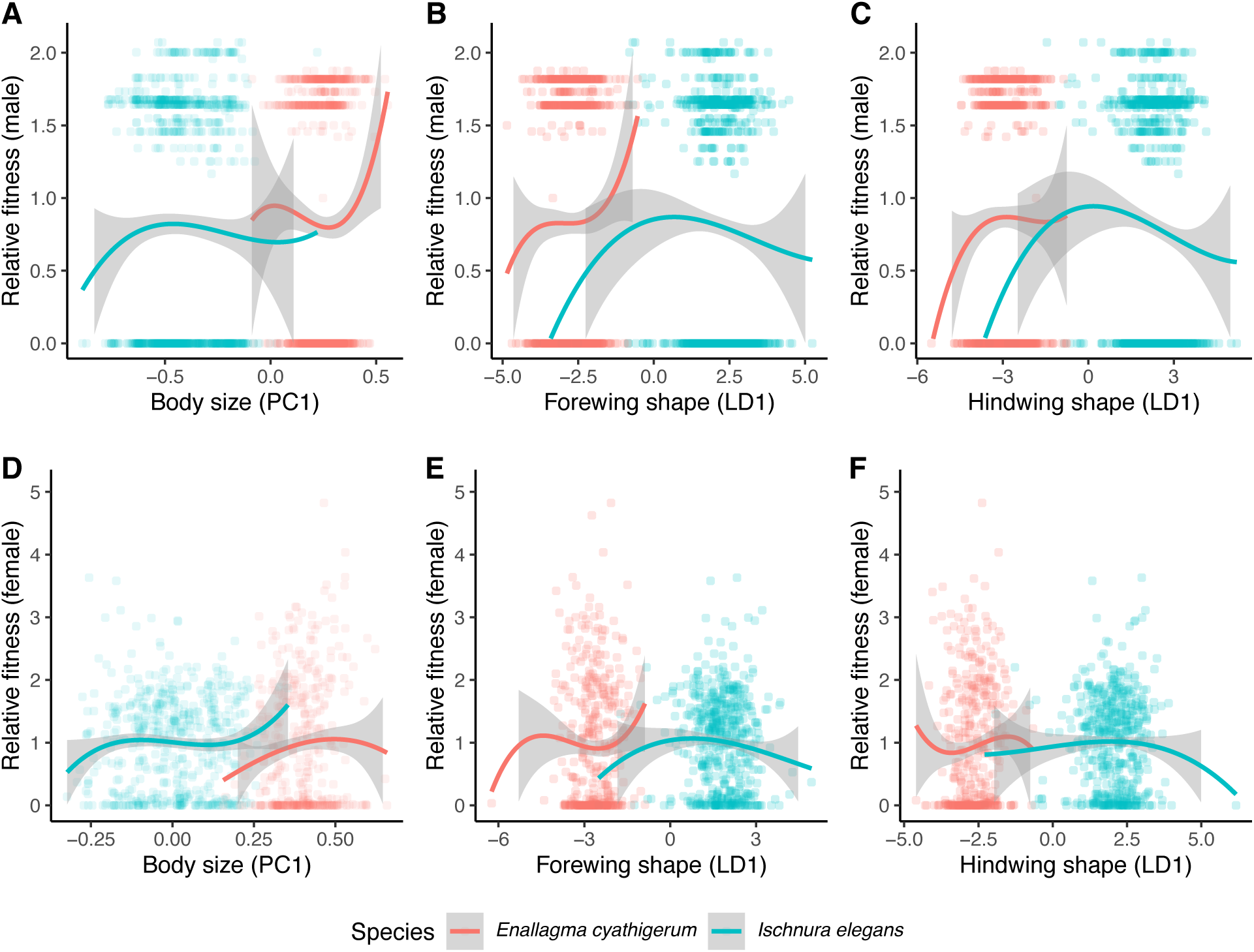
Univariate sexual selection (males, A-C) and fecundity selection (females, D-F) on body size (PC1) and wing shape (LD1) in *E. cyathigerum* and *I. elegans.* Fitness functions were visualized using cubic splines. Fitness data on the Y-axis were relativized at each population. The gray shaded regions around each spline represents the 95% confidence intervals. A) Selection on body size (PC1) in males, B) Selection on male forewing shape (LD1), C) Selection on male hindwing shape (LD1), D) Selection on body size (PC1) in females, E) Selection on female forewing shape (LD1) F) Selection on female forewing wing shape (LD1). The estimates for the coefficients of selection are presented in Table 1.

**Table 1:** Mean-standardized univariate linear (β) and quadratic (γ) selection gradients for body size (PC1), and wing shape (LD1) to complement Figure 4. Estimates within square brackets “[]” are variance-standardized selection gradients (traits are divided by the standard deviation of corresponding trait of the species before analysis). Estimates that are statistically significantly different from 0 at a significance level of p < 0.05 are shown in **bold**. All estimates are obtained from mixed-effect models that include sampled location as a random effect.

| Trait | Sex | $\beta \pm \text{SE}$ | | $\gamma \pm \text{SE}$ | |
| --- | --- | --- | --- | --- | --- |
|  |  | *I. elegans* | *E. cyathigerum* | *I. elegans* | *E. cyathigerum* |
| Body size PC1 | ♂ | 0.077±0.184 | -0.014±0.334 | -2.311±1.863 | **9.226±4.639<sup>(\*)</sup>** |
|  |  | [0.012±0.029] | [-0.001±0.031] | [-0.059±0.047] | [**0.078±0.039<sup>(\*)</sup>**] |
|  | ♀ | 0.172±0.251 | 1.103±0.700 | -0.770±3.322 | -17.758±11.540 |
|  |  | [0.018±0.022] | [0.012±0.022] | [0.005±0.022] | [-0.004±0.003] |
| Wing shape LD1 (forewing) | ♂ | -0.024±0.033 | 0.066±0.049 | **-0.052±0.025<sup>(\*)</sup>** | 0.109±0.093 |
|  |  | [-0.014±0.022] | [0.034±0.025] | [-0.027±0.018] | [0.007±0.015] |
|  | ♀ | -0.048±0.039 | -0.019±0.089 | -0.050±0.034 | 0.116±0.130 |
|  |  | [-0.027±0.027] | [-0.007±0.033] | [-0.033±0.028] | [0.002±0.010] |
| Wing shape LD1 (hindwing) | ♂ | -0.051±0.033 | 0.034±0.046 | **-0.050±0.023<sup>(\*)</sup>** | -0.124±0.074 |
|  |  | [-0.034±0.021] | [0.014±0.027] | [-0.010±0.012] | [-0.020±0.028] |
|  | ♀ | -0.006±0.040 | 0.072±0.097 | -0.034±0.028 | 0.056±0.199 |
|  |  | [-0.052±0.042] | [0.0001±0.003] | [-0.002±0.017] | [0.0002±0.0001] |

Note: \* indicates $0.01 < p < 0.05$ , \*\* indicates $0.001 < p < 0.01$ , \*\*\* indicates $p < 0.001$ .

Directional selection on wing shape LD1 was weak in both sexes and species (Fig. 2, Table 1). Estimates of β showed that selection tends to favor narrow and elongated wings in *I. elegans* while broad and round wings are favored in *E. cyathigerum*. Considering the mean shape differences between the two species (Fig. 1), these estimates suggest that, if all else being equal, directional selection would drive the wing shape to become progressively similar. The quadratic selection gradients (γ) suggested stabilizing selection (i.e., negative γ) on wing shape LD1 in all trait-sex-species combination except for forewing of *E. cyathigerum,* including estimates in males that were significantly different from zero (Table 1). Together with the result of β that favors intermediate wing shape of the two species, these results suggest that wing shape appear to be under weak stabilizing selection in both *I. elegans* and *E. cyathigerum*. For both size PC1 and wing shape LD1, variance-standardized gradients showed qualitatively equivalent results (Table 1). Results of correlational selection between wing shape and body size are presented in Supplementary Material (Table S1, Fig. S7, S8).

### Comparison of adaptive landscape between species

Comparisons of the adaptive landscape between *I. elegans* and *E. cyathigerum* are summarized in Table 2. Pair-wise comparisons between a model and its nested model based on AICc values indicate that the adaptive landscape is divergent between the two species in terms of the quadratic terms of body size in males (*Δ*AICc = 6.6 in a model with forewing, *Δ*AICc = 3.3 in a model with hindwing) and in terms of the linear and quadratic terms of body size in females (*Δ*AICc = 5.6 in a model with forewing, *Δ*AICc = 7.0 in a model with hindwing). Combining these results with estimates presented in Table 1, these results indicate that the disruptive selection on male body size in *E. cyathigerum* is absent or even takes the form of stabilizing selection in *I. elegans* (Table1, Fig. 2A) while in females, directional selection favoring large body sizes is stronger in *E. cyathigerum* than in *I. elegans*, and stabilizing selection appear to be slightly more prominent in *E. cyathigerum* than in *I. elegans* (Table1, Fig. 2D).

**Table 2.**
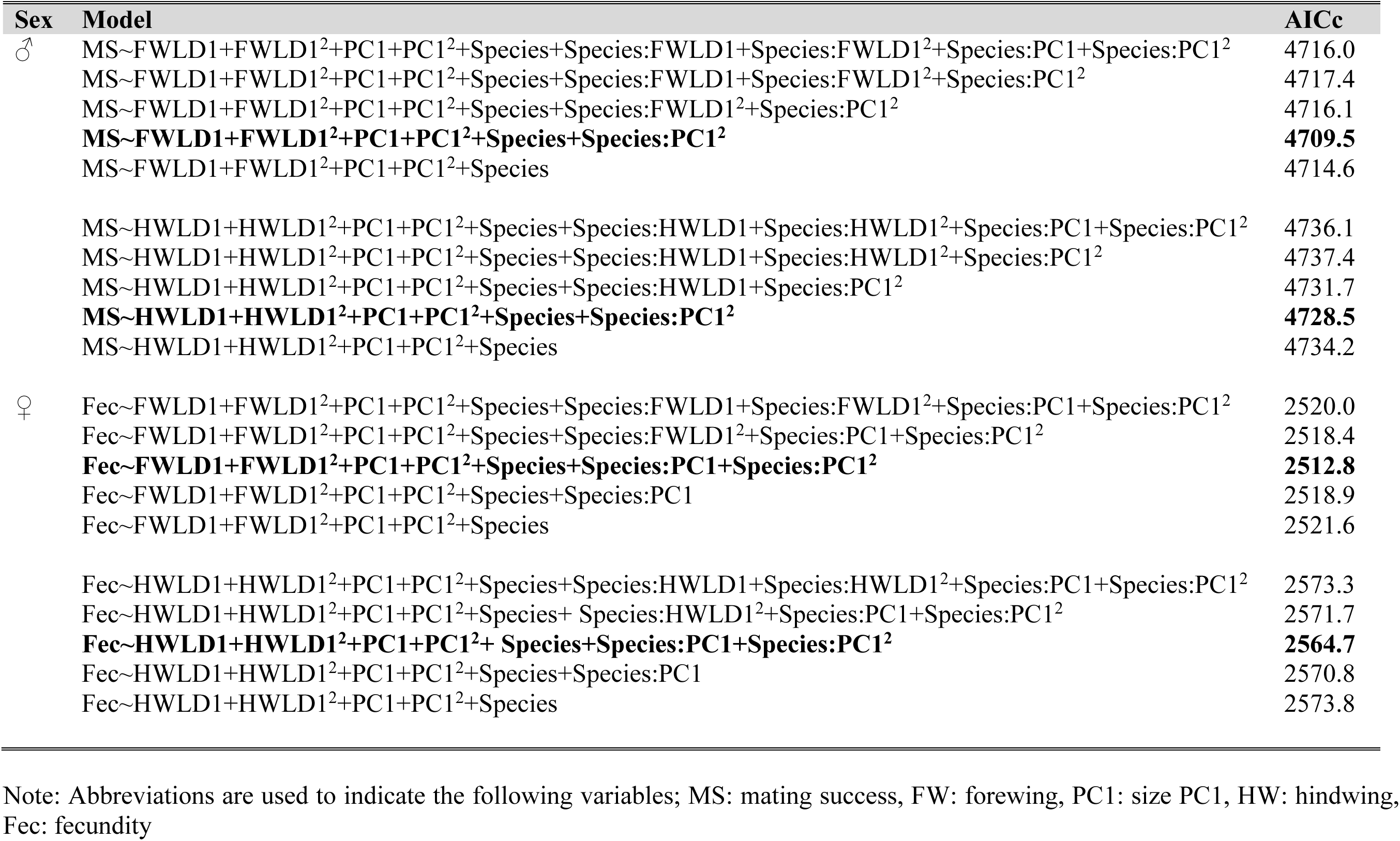
Model selection evaluating the difference of selection gradients between species. A full list of variables in a model and associated AICc values are presented. Note that the location of capture was included as a random effect in all models considered. The best model (i.e. a model with the lowest AICc) within a nested models are shown in **bold**.

### Selection gradients at population levels and sexual selection regime

Selection gradients evaluated at 17 populations with unique species-locale-year identity are presented in Table S2-S5. By comparing linear (β) and quadratic (γ) selection gradients at these population between *I. elegans* and *E. cyathigerum*, we found no statistical evidence that the local fitness surfaces are different between the two species (Fig. 3, Table S6). β on body size was far more variable than β on wing shape (variance of β: male body size = 2.36, female body size = 10.1, male forewing shape LD1 = 0.062, female forewing shape LD1 = 0.19, male hindwing shape LD1 = 0.021, female hindwing shape LD1 = 0.027, unit: (relative fitness/log_e_(mm))^2^ for body size, (relative fitness/centroid size)^2^ for wing shape), which globally fits with selection estimates at the species level that the adaptive landscape of wing shape was similar between the two species whereas that of body size showed more variation. Because γ was poorly estimated in most populations due to small sample sizes (Table S2-S6), rest of analyses will be performed on β.

**Figure 3.**
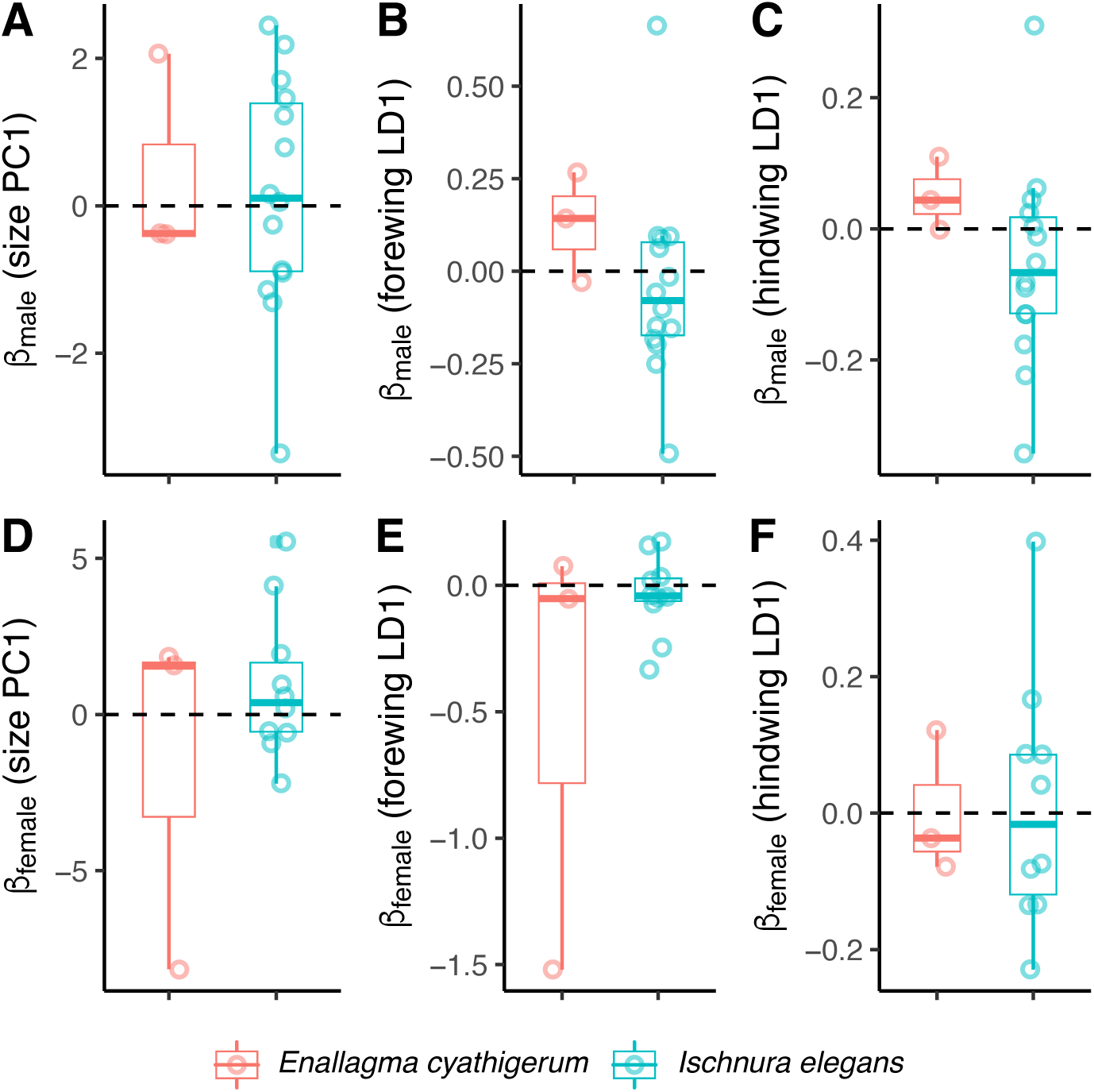
Boxplots (barline in the boxplot represent 50^th^ percentile and the intervals represent the range between 1.5 times above 75^th^ percentile and below 25^th^ percentile) comparing linear selection gradients (β) between *E. cyathigerum* and *I. elegans* evaluated at 17 populations. β of A) male body size, B) male forewing shape, C) male hindwing shape, D) female body size, E) female forewing shape, F) female hindwing shape are compared. The estimates for the mean and standard errors of these parameters as well as the statistical analysis for comparison between the two species (none are statistically different between the species) is presented in Table S6.

The summary of models relating β with matin system characteristics estimated at corresponding location and year are shown in Figure 4, Table S7 and Table S8. We found that the strength of directional selection on male body size is correlated with operational sex ratio (slope ± SE = 0.61 ± 0.15, p < 0.01) and opportunity for sexual selection (slope ± SE = 0.14 ± 0.05, p = 0.025), and that a non-significant trend exists with respect to androchrome frequency (slope ± SE = -4,43 ± 2.57, p = 0.11). These relationships (Fig. 4) suggest that large male body sizes are favored in populations where operational sex ratio is male-based, opportunity for sexual selection is high, and androchrome females (male-mimicking morph) are rare. Visual inspection of these relationships in *I. elegans* and *E. cyathigerum* as well as the lack of statistical support for the interaction term between species and mating system characteristics (Table S7, S8) suggest that the local sexual selection regime influences the strength of directional selection in male body size in a similar way in these two species. In addition, β on wing shape was related to local sexual selection regime in terms of OSR, androchrome frequency, and opportunity for selection (Table S8 and Figure S9). Although these relationships were somewhat weaker than those on β of body size, the way by which mating system characteristics are related to the strength of selection was remarkably consistent between *I. elegans* and *E. cyathigerum*. Overall, these results suggest that the direction and magnitude of sexual selection in these two species are governed by similar ecological processes.

**Figure 4:**
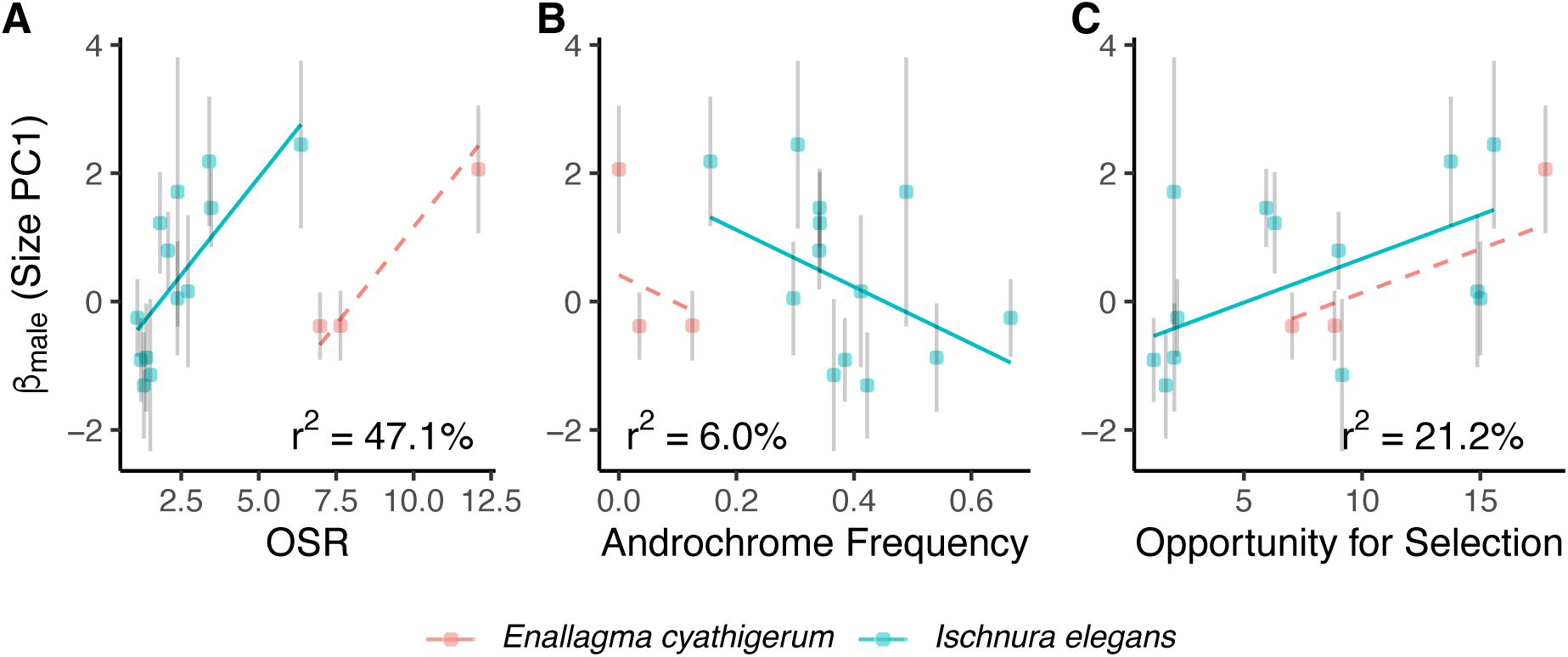
Relationship between directional selection gradient on male body size and (A) operational sex ratio (OSR), (B) androchrome frequency, (C) opportunity for selection. Each point represents a population with unique species-locale-year identity and error bar shows standard errors of selection gradients. Regression lines are derived from a weighted regression model (shown in Table S2-S5). Note that one population (*I. elegans* from Hoje A7 in 2021, see Table S2 for estimates of this population) was removed from plots for better visualization, while the regressions are from analyses using the whole dataset.

## Discussion

Research on phenotypic selection in natural populations has been a popular and largely successful endeavor (Endler, 1986, Mitchell-Olds & Shaw, 1987, Wade & Kalisz, 1990, Kingsolver et al., 2001), but its relevance to macroevolutionary dynamics remains poorly understood. In this study, we investigated how selection that varies in contemporary populations result in a stable adaptive zone, as has often been assumed in macroevolutionary models of phenotypic evolution (Hansen, 1997, Uyeda & Harmon, 2014, Ingram & Mahler, 2013). By estimating linear and quadratic selection on two fitness-related traits (body size and wing shape), we demonstrated that the adaptive landscape of wing shape is statistically indistinguishable between *I. elegans* and *E. cyathigerum* in terms of the form and intensity of selection. In contrast, the adaptive landscape of body size was more divergent between these two ecologically similar species. In addition, by analyzing the variation in selection at population level, we further showed that spatiotemporal variation in sexual selection on male body size arise from differences in mating system, and that local mating system influenced the direction and intensity of selection in a remarkably similar fashion in the two species. Below, we first propose graphical models that helps interpreting our results to build the link between contemporary selection and adaptive zone, before we discuss each finding in light of these models.

### A graphical model linking varying selection with macroevolution

To elucidate how selection that varies at the local level may result in a stable adaptive zone, let us consider three graphical models illustrating three plausible, not mutually exclusive, scenarios (Fig. 5). In the first model (Fig. 5A), the adaptive zone and the local adaptive landscape correspond to each other, and populations of each species simply oscillate around a shared adaptive peak. In the second model, the adaptive zone and landscape are corresponding, but two species occupy different parts of the adaptive zone (Fig. 5B). In this model, it is assumed that species-specific mean values are kept divergent, despite being constantly influenced by net directional selection favoring a shared adaptive peak, due to unmeasured variables that are sufficient to keep them apart but insufficient to move them away from the general selective regime imposed by their mode of life. Finally, in the third model, two species are subject to their own adaptive landscape with divergent adaptive peaks, while their selective regimes globally share ecological underpinnings. When environment changes, mean phenotypes of populations within each species evolve around respective divergent optima, but in a synchronous fashion such that they will never evolve away to form separate adaptive zones. In what follows, we refer to these models as Model A, B, and C. These models provide following predictions when two species share the adaptive zone: (1) When phenotypes are similar in the two species, selection gradients should be stabilizing in both species (Fig. 5A). (2) When phenotypes are dissimilar, the directional selection gradients should point to the same global phenotypic optima (Fig. 5B). (3) When phenotypes are dissimilar between species and multiple peaks are observed, local selection and ecological agents of selection should be correlated in a similar manner between the species (Fig. 5C).

**Figure 5:**
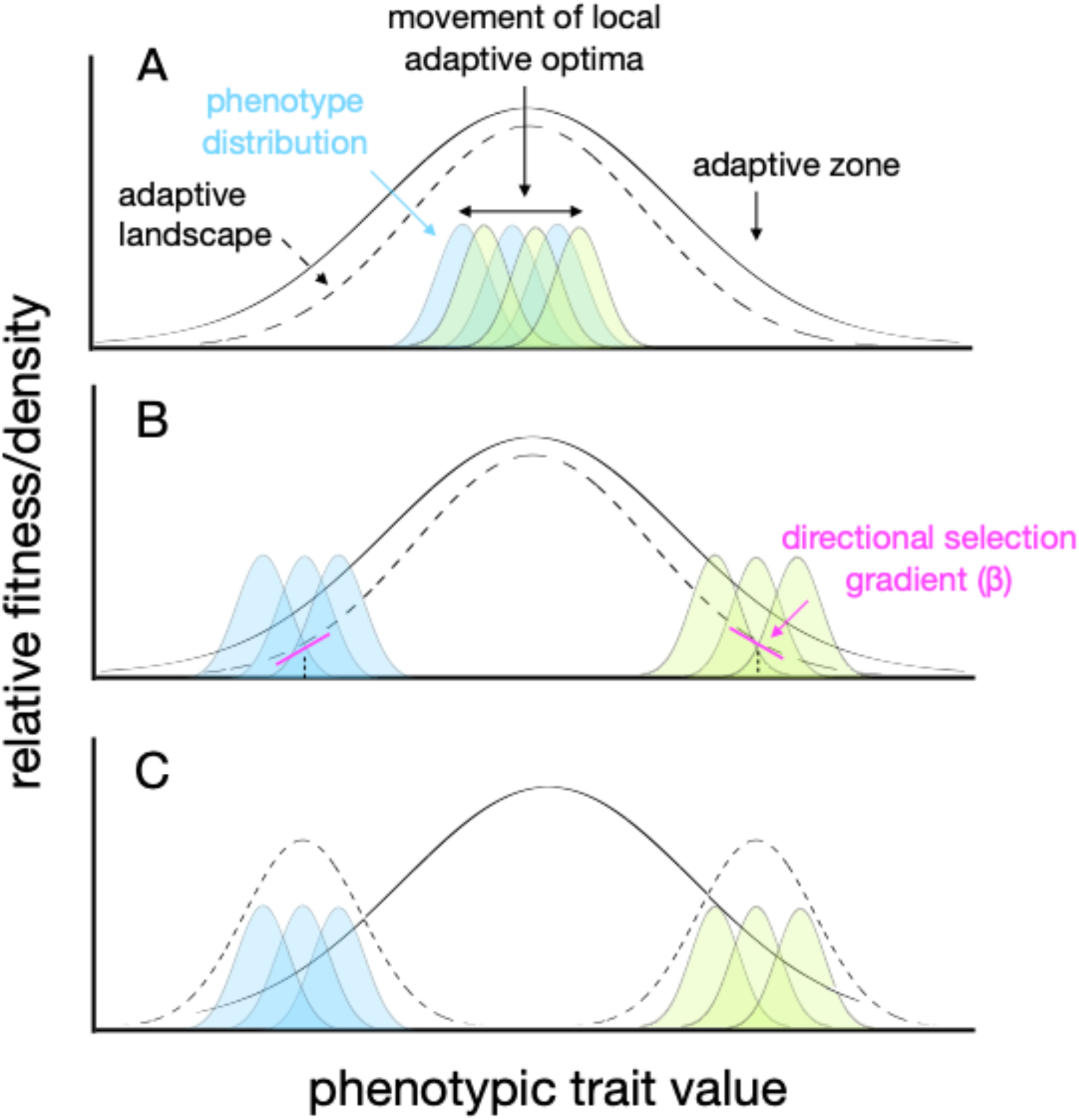
Three graphical models explaining how fluctuating selection can yield stable adaptive zone. Solid gaussian curve represents the adaptive zone occupied by populations of multiple species. Dashed gaussian curve represents the adaptive landscape of one or more species, and colored gaussian curve represents distribution of phenotypes within populations (two species are presented in different colors). The derivative of the adaptive landscape at the population mean is the directional selection gradient (β). (A) describes a case where the adaptive zone and the adaptive landscapes of two species correspond to each other. (B) describes a case where two species reside at different parts of a shared adaptive landscape that matches the adaptive zone. (C) describes a case where each species occupies their own adaptive landscape but are still part of a globally shared adaptive zone. Each model leads to specific predictions about the empirical estimates of β and γ as explained in the main text.

### Wing shape

Estimates of linear and quadratic selection on wing shape in *I. elegans* and *E. cyathigerum* revealed that wing shape is under a weak stabilizing selection (Fig. 2, Table 1). From a functional point of view, the signature of stabilizing selection itself may not be surprising because wing shape influences flight performance, which is the key determinant of virtually all fitness-related activities of adult damselflies including feeding, territorial defense, mating or predator avoidance (Corbet, 1999). What surprised us is that, despite the deep divergence between the two species (> 12 million years; Willink et al., 2024b, Blow et al., 2021), there was no evidence that the form and intensity of selection differ between them. One interpretation of this finding is that wing shape is maintained around two species-specific adaptive peaks that are similar in form but are part of two separate adaptive zones. Alternatively, these two species may occupy different parts of a shared adaptive zone (Fig. 5B). Our results are in favor of the latter, because estimates of directional selection (β) revealed that, among eight estimates, all β on wing shape in *I. elegans* were negative in sign, whereas β on wing shape in *E. cyathigerum* were all positive in sign except for female forewing (Table 1).

By visualizing the population-specific selection gradients of wing shape together with the global selective surfaces (Fig. 6), we further found that selection often favors a wing shape intermediate between these two species, with a possible exception in hindwing of females where β seems to push populations to species-specific adaptive peaks (Fig. 6D). It is conceivable that the intermediate wing phenotype represents a global adaptive optimum determined by some fundamental mechanical and physiological demands of flight. Each one of these two species may occupy a fraction of this adaptive zone according to their ecological mode of life, that are globally similar but exhibit differences reflecting their subtle ecological differences (Fig. S11, Supplementary Result and Discussion). Together with the result that wing shape of both species are generally under stabilizing form of selection, the adaptive landscapes of wing shape are likely yield a stable adaptive zone through a combination of Model A and B. By extending the examination to 5 PC axes of shape (cumulatively explain 82.2% of all wing shape variation), we confirmed that our interpretation generally applies to all five shape dimensions (Fig. S10).

**Figure 6.**
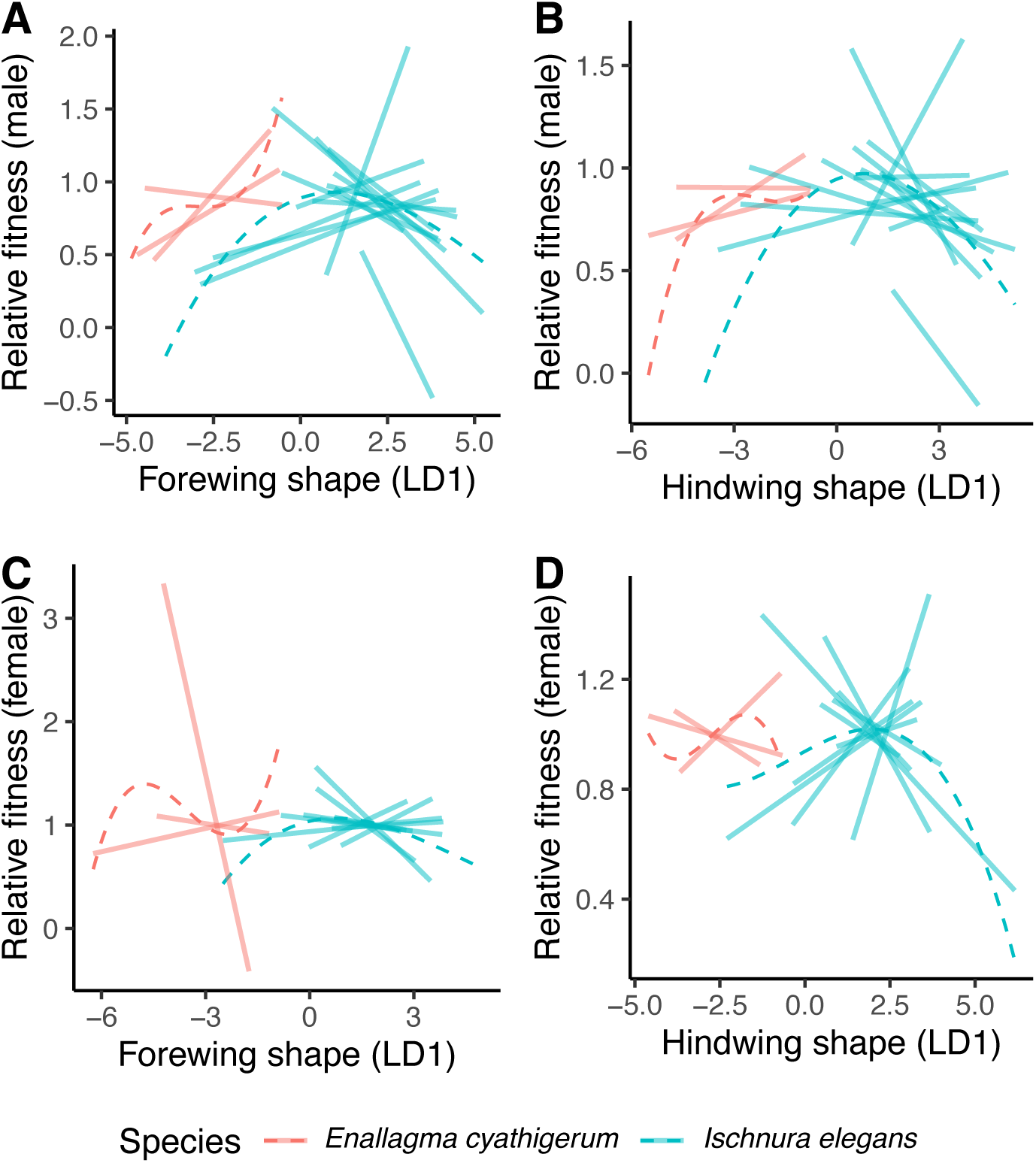
Comparison of species-specific fitness surface with directional selection gradients at each locality (3 populations in *E. cyathigerum* and 14 populations in *I. elegans*). Sexual selection (males, A-B) and fecundity selection (females, C-D) on wing shape (LD1, both fore- and hindwings) are plotted. Fitness functions of each species (dashed lines) were visualized using cubic splines. Fitness data on the Y-axis were relativized at each location. Directional selection gradients (solid lines) were estimated using a regression model of relative fitness against LD1 at each location. Estimates of these models are presented in Table S2-S5.

Our analyses illustrate that multiple species may occupy a single adaptive zone even when individual selection estimates at any given place and time may give an impression of directional selection (Svensson & Sinervo, 2004, Haller & Hendry, 2014). Recent evidence suggests that the stabilizing form of adaptive landscape can be maintained by spatiotemporally varying selection (Beausoleil et al., 2023, Stroud et al., 2023, de Villemereuil et al., 2020). Our study adds another empirical evidence from natural populations supporting this view.

### Body size

Compared to selection on wing shape, selection on body size was considerably more variable. Females in both species experienced positive directional selection towards large body size, and this fecundity selection was stronger in magnitude in *E. cyathigerum* than in *I. elegans* (Fig. 3, Table 1, Table 2). These two species differ slightly in terms of the strength of stabilizing selection (Table 2) but there is large uncertainty of these estimates. Visual inspections of the data suggests that the stabilizing selection on female body size is weak in both species. An elevated sexual conflict through intense male mating harassment in *I. elegans* compared to *E. cyathigerum* (Fig. S11) can potentially explain divergence in selection on female body size. Male mating harassment in *I. elegans* has been shown to be mainly directed towards high-fecundity females (Gosden & Svensson, 2009). In other insects such as *Drosophila*, large females have intrinsically higher fecundity, but they also suffer more from male mating attempts that reduce these intrinsic fitness advantages of large females (Long et al., 2009, Chenoweth et al., 2015). In contrast to the situation in *I. elegans*, in *E. cyathigerum,* benefits of large size in females might be less affected by male mating harassment because male mating harassment appear to be lower in *E. cyathigerum* compared to *I. elegans* based on several mating system indices (Supplementary Results, Table S9, Fig. S11).

For example, the frequency of androchrome females in *E. cyathigerum* is substantially lower than in *I. elegans*. Previous studies have shown that androchrome females in damselflies benefit from reduced male mating harassment in the genus *Ischnura* (Robertson, 1985, Cordero et al., 1998, Gosden & Svensson, 2009, Blow et al., 2021). The lower frequency of androchrome females in *E. cyathigerum* compared to *I. elegans* might therefore reflect relaxed mating harassment and reduced sexual conflict in the former species. In addition, female *E. cyathigerum* are more locally segregated from conspecific males of the same species, as previously documented in other species of this genus (Stoks, 2001, Steele et al., 2011), whereas in *I. elegans*, males and females typically co-occupy the same microhabitat and sex ratios are more balanced in this species (Fig. S11). Sexual habitat segregation might therefore further rescue females of *E. cyathigerum* from the cost of male mating harassment, enabling them to capitalize on greater fecundity advantages of large body size compared to *I. elegans*.

Overall, we interpret these results as indicating that selection on female body size in *E. cyathigerum* and *I. elegans* are governed by similar ecological factors that differ in their intensity. As circumstantial evidence for this, *E. cyathigerum* is significantly bigger than *I. elegans* in body size (Fig. 2). This may be the consequence of a strong directional selection in *E. cyathigerum* within an adaptive zone that they share with *I. elegans*. We suggest that contemporary selection in female body size of the studied damselflies results in a stable adaptive zone through a combination of Model B and C.

In males, we found divergent patterns of selection on body size between the two species (Fig. 2A). *I. elegans* is characterized by a weak (non-significant) stabilizing selection while *E. cyathigerum* experienced disruptive selection (Fig. 2A). These results indicate that sexual selection on male body size is highly context-dependent in these damselflies (see Supplementary Result for related discussion on correlational selection). Previous studies have shown that the form and magnitude of sexual selection in damselflies can vary in a fine spatial and temporal scale depending on local social structure, such as the density of female color morphs (Gosden & Svensson, 2008) or phenotypic distribution of male and female body sizes (Steele et al., 2011). These results indicate either that male body sizes are evolving on a complex multipeak adaptive zones that separate the two species over macroevolutionary time scales, like the case of *Anolis* lizard (Stroud et al., 2023) and *Geospiza* finch (Beausoleil et al., 2023), or, alternatively that shared ecological agents keeps male body sizes apart from each other within a shared adaptive zone (Model C).

### Ecological causes of variation in sexual selection

By incorporating mating system variables from different study sites into our analyses, we sought to explore the validity of Model C in explaining the variation in sexual selection on male body size. We found that selection on male body size became stronger when the operational sex ratio (OSR) was male-biased (Fig. 4A), when androchrome females were rare (Fig. 4B) and when the opportunity for selection was high (Fig. 4C). In insects that engage in scrambling male-male competition, OSR and the opportunity for selection are important descriptors of mating systems. Such parameters have previously shown to be correlated with sexual selection regimes in water striders (Arnqvist, 1992) and ambush bugs (Punzalan et al., 2010). Our findings that low androchrome frequency seem to strengthen directional selection on male body size mirrors previous result of Gosden and Svensson (2008) who demonstrated a similar pattern in *I. elegans* based on extensive spatiotemporal replicates (60 populations over 5 years) compared to ours (17 populations over 2 years). One explanation for this is that in populations where androchrome females are common, males might also suffer from male mating attempts due to mistaken mate recognition, as previously shown in *I. ramburii* (Gering, 2017). Consequently, in populations with high frequencies of androchrome females, smaller males might be favored to reduce similarity with androchrome females and thereby also reduce male-male interference.

Importantly, the local mating environments affected the magnitude of directional sexual selection in *I. elegans* and *E. cyathigerum* in a remarkably similar fashion (Fig. 4, Fig. S9). This result suggests that sexual selection regimes in males of these two species are shaped by the same factors governing their mating systems, which should partly reflect the resemblance of ecological niche among pond damselflies (McPeek & Brown, 2000, Siepielski et al., 2010). This indicates that the difference in the adaptive landscapes of male body size between *I. elegans* and *E. cyathigerum* likely reflect slight differences in mating systems (Supplementary Results, Table S9, Fig. S11). Furthermore, this also indicates that selection regimes of the two studied pond damselflies should vary in a synchronous manner whenever variables governing their mating systems change in nature. Over macroevolutionary time scales, such synchronous dynamics of adaptive landscapes with closely located but distinct optima will cause the populations of each species to perpetually oscillate around their adaptive peaks, while still not moving away too far from each other. If the ecological niche—here we refer to a suite of ecological and environmental circumstances that set the conditions for the existence of organisms—shared by residing species and populations are persistent over macroevolutionary timescales, a stable adaptive zone will form (Simpson, 1944). A plausible mechanism that stabilizes sexual selection dynamics in pond damselflies is the negative frequency-dependent selection driven by the frequency of female-limited color polymorphisms (Takahashi et al., 2010, Le Rouzic et al., 2015, Svensson et al., 2005). Altogether, our results support Model C, where the distinct adaptive landscapes between *I. elegans* and *E. cyathigerum* in male body size are part of a single adaptive zone representing an ecological mode of life shared by pond damselflies (McPeek & Brown, 2000, Siepielski et al., 2010).

## Conclusion

The adaptive landscape has been argued to provide a conceptual bridge between micro- and macroevolution for more than two decades (Arnold et al., 2001), but its application had primarily been conceptual and metaphorical until recently (Beausoleil et al., 2023, Stroud et al., 2023, Martin, 2016). Our study extends ongoing attempts to incorporate insights from phenotypic selection research into models focused on adaptive radiation and macroevolution (Arnold, 2023, Svensson, 2023, Pennell & Jiang, 2024). The demonstration of ecological correlates of variation in selection suggests that a better understanding of the ecological causes of selection (Svensson & Sinervo, 2000, MacColl, 2011, Opedal, 2021) hold great promise to explore the link between micro- and macroevolution. While the two-species approach in this study is limited in its scope (but see Price, 1997 for arguments in favor of two-species comparative studies), it does not affect our points to stimulate future empirical study on the micro-macro link within the conceptual framework of the adaptive landscape (Arnold, 2023, Svensson & Calsbeek, 2012, Hansen et al., 2023). Phenotypic selection studies that are replicated across different species that covers a wide range of phenotype space and ecological circumstances provides a promising way forward to consolidate the emerging bridge between micro- and macroevolution (Tsuboi et al., 2024, Rolland et al., 2023).

## Acknowledgements

We thank Stephen de Lisle and Viktor Nilsson-Örtman for discussions on earlier versions of analyses and manuscripts, Seth Donoughe for helping us with image processing, and Benjamin Jarrett, Emma Kärrnäs, Moritz Lürig, Ayushi Mahajan, Muskaan, Sofie Nilén, Karolina Pehrson, Moa Metz and Kajsa Svensson for assisting our field work. We also thank constructive criticisms from Gabrielle Marroig, Michel Morrissey, and David Punzalan that substantially improved this manuscript. This study was supported by the Swedish Research Council grant for EIS (2020-03123) and MT (2016-06635).

## Data accessibility

Data and code will be archived in GitHub upon acceptance.

