## Supplementary Material for "Persistence of the ecological niche in pond damselflies underlies a stable adaptive zone despite varying selection"

### §1 Supplementary Methods

**Comparison of social organization indices between *Ischnura elegans* and *Enallagma cyathigerum***: We tested for the difference in six social organization indices (see main text for details of those indices) between *E. cyathigerum* and *I. elegans* using a mixed effect model implemented in lme4 package with one of the six parameters as the response variable, species as the fixed explanatory variable and sampling year and location at which individuals were captured as the random effect.

### §2 Supplementary Results

**Social organization and mating system:** Comparison of social organization between *I. elegans* and *E. cyathigerum* are presented in Figure S11. We found statistical evidence for differences in social organization and mating system between the two species in all examined parameters except for male density (Table S9). Males of *I. elegans* were roughly 1.5 times as more often found as mating couple as *E. cyathigerum* (proportion of copula; *I. elegans*: mean ± SE = 0.133 ± 0.023, *E. cyathigerum*: 0.091 ± 0.018), experienced 33% less opportunity of sexual selection ($I_s$) than *E. cyathigerum* ($I_s$; *I. elegans*: mean ± SE = 9.76 ± 2.31, *E. cyathigerum*: 14.48 ± 2.13), and females of *I. elegans* had 1.6 times as high frequency of the male-mimicking phenotype (*androchrome*) as *E. cyathigerum* (androchrome frequency; *I. elegans*: mean ± SE = 0.46 ± 0.05, *E. cyathigerum*: 0.28 ± 0.12). The operational sex-ratio (OSR) is male-biased in both species, but significantly more so in *E. cyathigerum* (OSR; *I. elegans*: mean ± SE = 2.87 ± 0.85, *E. cyathigerum*: 8.72 ± 0.75), which partly reflects low overall density of females near the pond in this species compared to *I. elegans*.

**Correlational selection**: The multivariate selection gradients, including correlational selection between size PC1 and shape LD1, are presented in Table S1, Figure S7, and Figure S8. Estimates of gradients are generally consistent with those obtained from univariate models, with one notable exception. In males of *E. cyathigerum*, the evidence for disruptive selection on body size in the univariate estimate of $\gamma$ is shown to result from a combination of directional selection favoring small size ($\beta$ ± SE = -4.47 ± 1.79), a non-significant trend of disruptive selection on wing shape ($\gamma$ ± SE = 7.22 ± 4.76), and a marginal effect of negative correlational selection between body size and wing shape ($\gamma$ ± SE = -1.03 ± 0.56). Visualization of this selection surface revealed a complex adaptive topography in males of *E. cyathigerum* characterized by a fitness saddle with two fitness peaks where large and small males had higher mating success (Fig. S7). The fitness surface of males of *I. elegans* was flatter with a tendency for weak stabilizing selection on body size ($\gamma$ ± SE = -2.63 ± 2.05). By contrast, the adaptive landscapes of females were simpler than males in both species. Correlational selection was virtually absent, and fitness monotonically increased with body size consistent with the univariate estimates.

### §3 Supplementary Discussion

***Ischnura elegans and Enallagma cyathigerum* differ in their social organization**: Comparisons of mating system parameters in *I. elegans* and *E. cyathigerum* revealed interesting differences (Fig. S11). Compared to *E. cyathigerum*, *I. elegans* have higher female densities (Fig. S11B), more even operational sex ratio (Fig. S11C), higher frequency of androchrome females (Fig. S11D) and lower the opportunity for sexual selection (Fig. S11F). These species differences suggest that mating systems are different between *I. elegans* and *E. cyathigerum.* In *E. cyathigerum*, there is a classical mating system based on male-male competition over the opportunity for mating. In contrast, in *I. elegans*, sexual conflict through male mating harassment appears to be the primary mechanism of sexual selection (Fig. S11C). A potentially important qualitative difference between the mating system of these species might be differences in postcopulatory mate guarding during female oviposition (Corbet, 1999). In *I. elegans*, females oviposit alone, unguarded by the male, where they are likely to be more susceptible to mating harassment from other males who try to mate with vulnerable females during their oviposition phase. In contrast, in *E. cyathigerum*, males remain attached to the female in tandem position, even during female oviposition, suggesting that females are less vulnerable to male mating harassment (personal observations; Fig. S1E). These subtle but fundamental differences in pre- and post-mating behavior are likely causes of mating system differences and social structure between these two ecologically similar species.

**Weak correlational selection between body size and wing shape**: We found that none of correlational selection were statistically supported (Table S1). Yet there was one notable difference between univariate and multivariate selection on body size in males. In univariate model, we found that *I. elegans* is characterized by a weak (non-significant) stabilizing selection while *E. cyathigerum* is under a disruptive selection (Table 1). When correlational selection between wing shape and body size were considered, the disruptive selection in *E. cyathigerum* was instead described by a combination of negative directional selection towards small body size and a negative correlational selection between body size and wing shape (Table S1). These results suggest that the pattern of sexual selection in males of *E. cyathigerum* is not just context dependent as we argued in the main text, but also influenced by indirect effect of sexual selection acting on wing shape.

Table S1: Mean-standardized linear (β) and quadratic (γ) selection gradients of body size (PC1), and wing shape (LD1) that characterizes multivariate selection surfaces shown in Figure S7 and S8. Estimates that are statistically significantly different from 0 at a significance level of $p < 0.05$ are shown in **bold**. All estimates are obtained from mixed-effect models that include sampled location as a random effect.

| Sex | Parameters | estimates ± SE (forewing) | | estimates ± SE (hindwing) | |
|---|---|---|---|---|---|
| | | *I. elegans* | *E. cyathigerum* | *I. elegans* | *E. cyathigerum* |
| ♂ | Wing shape LD1 | -0.010±0.076 | 0.513±0.271 | -0.003±0.073 | -0.223±0.277 |
| | Wing shape LD1$^2$ | **-0.071±0.034**$^{(*)}$ | 0.072±0.095 | -0.053±0.030 | -0.135±0.075 |
| | Body size PC1 | -0.425±0.833 | **-4.474±1.791**$^{(*)}$ | -0.666±0.869 | **-3.615±1.805**$^{(*)}$ |
| | Body size PC1$^2$ | -2.633±2.053 | 7.222±4.761 | -2.244±1.960 | 8.393±4.666 |
| | LD1 × PC1 | -0.246±0.261 | -1.034±0.563 | -0.076±0.246 | -0.584±0.527 |
| ♀ | Wing shape LD1 | 0.042±0.073 | 0.043±0.561 | 0.093±0.066 | 0.179±0.623 |
| | Wing shape LD1$^2$ | -0.062±0.042 | 0.122±0.129 | -0.049±0.31 | 0.070±0.210 |
| | Body size PC1 | 0.389±0.757 | 9.411±4.847 | 1.196±0.807 | 9.147±5.132 |
| | Body size PC1$^2$ | -0.026±3.714 | -15.694±11.704 | -2.231±3.646 | -17.826±12.028 |
| | LD1 × PC1 | -0.167±0.368 | 0.745±1.128 | -0.459±0.350 | 0.285±1.286 |

Note: * indicates $0.01 < p < 0.05$, ** indicates $0.001 < P < 0.01$, *** indicates $p < 0.001$.

Table S2. Summary of selection gradients in body size PC1 of *I. elegans* based on mating success (male) and numbers of eggs laid (female) evaluated at each population and year. Mean-standardized linear (β) and quadratic (γ) selection gradients and associated standard errors (SE) are estimated from linear regression models with fitness component (relativized at each population) against body size PC1. Estimates that are statistically significantly different from 0 at a significance level of $p < 0.05$ are shown in **bold**. N refers to sample size, Mean and SD refer to mean and standard deviation of body size PC1 (centered at the global mean, unit: $\log_e$(mm)) at each population.

| Sex | Year | Locale | N | Mean | SD | β ± SE | γ ± SE |
|---|---|---|---|---|---|---|---|
| ♂ | 2020 | Borgeby | 28 | -0.456 | 0.154 | 0.16±1.19 | -13.3±15.9 |
| | | Bunkeflostrand | 36 | -0.596 | 0.121 | -1.15±1.19 | -15.7±12.8 |
| | | Flyinge 30 A3 | 50 | -0.424 | 0.105 | **2.18±1.01*** | **35.7±16.1*** |
| | | Generp | 80 | -0.450 | 0.131 | **1.46±0.61*** | 9.34±6.67 |
| | | Hoje A14 | 96 | -0.447 | 0.101 | -0.87±0.84 | 8.65±11.6 |
| | | Hoje A7 | 71 | -0.370 | 0.163 | 0.79±0.60 | -2.50±6.56 |
| | | Ilstorp | 55 | -0.528 | 0.141 | 1.22±0.79 | -6.94±9.00 |
| | | Vombs Vattenverk | 136 | -0.466 | 0.087 | -1.31±0.83 | 22.0±13.1 |
| | 2021 | Borgeby | 33 | -0.389 | 0.142 | 2.45±1.30 | -9.06±13.9 |
| | | Flyinge 30 A3 | 24 | -0.384 | 0.163 | 0.049±0.89 | -18.6±11.0 |
| | | Generp | 21 | -0.240 | 0.063 | 1.71±2.10 | 49.9±63.4 |
| | | Hoje A7 | 21 | -0.198 | 0.047 | -3.36±3.87 | 132.0±155.0 |
| | | Ilstorp | 56 | -0.408 | 0.188 | -0.25±0.61 | -7.10±8.29 |
| | | Vombs Vattenverk | 102 | -0.306 | 0.152 | -0.91±0.65 | -4.14±6.09 |
| ♀ | 2020 | Flyinge 30 A3 | 25 | 0.0011 | 0.094 | **5.53±1.56**** | 23.2±35.2 |
| | | Generp | 45 | -0.057 | 0.132 | -2.22±1.11 | -26.3±15.7 |
| | | Hoje A14 | 50 | -0.022 | 0.078 | -0.90±1.33 | -13.9±25.4 |
| | | Hoje A7 | 37 | 0.022 | 0.118 | -0.54±1.34 | -20.8±21.6 |
| | | Ilstorp | 31 | -0.076 | 0.113 | -0.60±1.14 | 9.79±18.7 |
| | | Vombs Vattenverk | 67 | -0.038 | 0.107 | **1.92±0.71**** | -4.23±7.7 |
| | 2021 | Generp | 40 | 0.114 | 0.050 | 4.10±2.15 | -28.4±58.0 |
| | | Hoje A14 | 22 | 0.033 | 0.130 | 0.98±0.76 | -8.43±9.83 |
| | | Ilstorp | 27 | -0.026 | 0.133 | 0.19±1.45 | -14.4±24.6 |
| | | Vombs Vattenverk | 34 | 0.038 | 0.125 | 0.57±0.87 | -4.55±13.2 |

Note: * indicates $0.01 < p < 0.05$, ** indicates $0.001 < P < 0.01$, *** indicates $p < 0.001$.

Table S3. Summary of selection gradients in forewing shape LD1 of *I. elegans* based on mating success (male) and numbers of eggs laid (female) evaluated at each population and year. Mean-standardized linear (β) and quadratic (γ) selection gradients and associated standard errors (SE) are estimated from linear regression models with fitness component (relativized at each population) against forewing shape LD1. Estimates that are statistically significantly different from 0 at a significance level of p < 0.05 are shown in **bold**. N refers to sample size, Mean and SD refer to mean and standard deviation of forewing shape LD1 (unit: centroid size) at each population.

| Sex | Year | Locale | N | Mean | SD | β ± SE | γ ± SE |
|---|---|---|---|---|---|---|---|
| ♂ | 2020 | Borgeby | 28 | 1.89 | 1.42 | 0.10±0.13 | 0.020±0.14 |
| | | Bunkeflostrand | 36 | 2.65 | 0.93 | -0.25±0.15 | -0.25±0.23 |
| | | Flyinge 30 A3 | 50 | 2.16 | 0.91 | -0.10±0.12 | 0.20±0.14 |
| | | Generp | 80 | 2.08 | 0.99 | 0.095±0.083 | -0.085±0.092 |
| | | Hoje A14 | 96 | 2.28 | 0.60 | -0.15±0.14 | -0.65±0.30 |
| | | Hoje A7 | 71 | 2.04 | 1.13 | 0.063±0.087 | 0.14±0.081 |
| | | Ilstorp | 55 | 2.58 | 0.80 | -0.058±0.14 | -0.062±0.27 |
| | | Vombs Vattenverk | 136 | 2.34 | 0.73 | -0.015±0.10 | -0.20±0.15 |
| | 2021 | Borgeby | 33 | 1.97 | 0.51 | 0.66±0.37 | -0.23±0.88 |
| | | Flyinge 30 A3 | 24 | 2.14 | 0.52 | -0.15±0.29 | 1.44±1.37 |
| | | Generp | 21 | 2.44 | 0.49 | -0.49±0.25 | 0.44±0.36 |
| | | Hoje A7 | 21 | 1.93 | 0.90 | 0.087±0.21 | 0.34±0.43 |
| | | Ilstorp | 56 | 2.25 | 0.66 | -0.18±0.17 | 0.26±0.52 |
| | | Vombs Vattenverk | 102 | 2.05 | 0.77 | -0.20±0.13 | -0.11±0.18 |
| ♀ | 2020 | Flyinge 30 A3 | 25 | 1.79 | 0.49 | -0.071±0.37 | 0.24±0.88 |
| | | Generp | 45 | 1.85 | 1.02 | 0.034±0.15 | **-0.26±0.12*** |
| | | Hoje A14 | 50 | 1.83 | 0.65 | -0.049±0.16 | -0.34±0.28 |
| | | Hoje A7 | 37 | 1.53 | 1.02 | -0.040±0.15 | -0.0034±0.12 |
| | | Ilstorp | 31 | 2.23 | 0.72 | 0.020±0.18 | 0.17±0.34 |
| | | Vombs Vattenverk | 67 | 1.86 | 0.71 | **-0.33±0.11**** | 0.35±0.21 |
| | 2021 | Generp | 40 | 1.63 | 0.61 | -0.25±0.18 | -0.31±0.36 |
| | | Hoje A14 | 22 | 1.66 | 0.77 | -0.044±0.13 | 0.18±0.37 |
| | | Ilstorp | 27 | 2.05 | 0.69 | 0.17±0.28 | -0.21±0.55 |
| | | Vombs Vattenverk | 34 | 1.34 | 0.75 | 0.16±0.14 | -0.055±0.27 |

Note: * indicates 0.01 < p < 0.05, ** indicates 0.001 < P < 0.01, *** indicates p < 0.001.

Table S4. Summary of selection gradients in hindwing shape LD1 of *I. elegans* based on mating success (male) and numbers of eggs laid (female) evaluated at each population and year. Mean-standardized linear (β) and quadratic (γ) selection gradients and associated standard errors (SE) are estimated from linear regression models with fitness component (relativized at each population) against hindwing shape LD1. Estimates that are statistically significantly different from 0 at a significance level of $p < 0.05$ are shown in **bold**. N refers to sample size, Mean and SD refer to mean and standard deviation of hindwing shape LD1 (unit: centroid size) at each population.

| Sex | Year | Locale | N | Mean | SD | β ± SE | γ ± SE |
|---|---|---|---|---|---|---|---|
| ♂ | 2020 | Borgeby | 28 | 2.09 | 1.56 | 0.045±0.12 | -0.079±0.075 |
| | | Bunkeflostrand | 36 | 2.62 | 0.75 | -0.18±0.19 | -0.13±0.38 |
| | | Flyinge 30 A3 | 51 | 2.18 | 0.89 | -0.090±0.12 | 0.044±0.083 |
| | | Generp | 80 | 2.24 | 1.05 | -0.012±0.079 | **-0.14±0.061*** |
| | | Hoje A14 | 96 | 2.39 | 0.65 | -0.082±0.13 | -0.046±0.27 |
| | | Hoje A7 | 71 | 2.12 | 1.21 | -0.052±0.082 | -0.011±0.059 |
| | | Ilstorp | 55 | 2.66 | 0.65 | 0.024±0.18 | 0.37±0.38 |
| | | Vombs Vattenverk | 136 | 2.51 | 0.69 | -0.13±0.11 | -0.0025±0.15 |
| | 2021 | Borgeby | 33 | 2.23 | 0.68 | 0.31±0.28 | -0.44±0.51 |
| | | Flyinge 30 A3 | 25 | 2.36 | 0.58 | 0.062±0.24 | 0.22±0.65 |
| | | Generp | 21 | 2.61 | 0.62 | -0.22±0.21 | 0.047±0.47 |
| | | Hoje A7 | 21 | 2.11 | 0.76 | -0.34±0.23 | 0.43±0.41 |
| | | Ilstorp | 56 | 2.43 | 0.77 | 0.0039±0.15 | 0.084±0.28 |
| | | Vombs Vattenverk | 101 | 2.27 | 0.71 | -0.13±0.14 | -0.088±0.29 |
| ♀ | 2020 | Flyinge 30 A3 | 25 | 2.13 | 0.61 | -0.13±0.30 | -1.42±1.11 |
| | | Generp | 45 | 2.07 | 1.00 | 0.087±0.15 | -0.15±0.12 |
| | | Hoje A14 | 50 | 2.00 | 0.58 | 0.042±0.18 | 0.47±0.12 |
| | | Hoje A7 | 37 | 1.95 | 1.23 | -0.14±0.13 | -0.053±0.055 |
| | | Ilstorp | 30 | 2.50 | 0.78 | -0.074±0.17 | 0.022±0.32 |
| | | Vombs Vattenverk | 67 | 2.12 | 0.65 | -0.23±0.12 | -0.13±0.30 |
| | 2021 | Generp | 40 | 1.80 | 0.65 | -0.081±0.17 | 0.19±0.32 |
| | | Hoje A14 | 22 | 1.76 | 0.77 | 0.086±0.13 | -0.21±0.20 |
| | | Ilstorp | 27 | 2.37 | 0.52 | 0.40±0.37 | 0.11±0.95 |
| | | Vombs Vattenverk | 33 | 1.62 | 0.75 | 0.17±0.15 | -0.086±0.27 |

Note: * indicates $0.01 < p < 0.05$, ** indicates $0.001 < P < 0.01$, *** indicates $p < 0.001$.

Table S5. Summary of selection of *E. cyathigerum* based on mating success (male) and numbers of eggs laid (female) evaluated at each population. All populations were sampled in 2021. Mean-standardized linear (β) and quadratic (γ) selection gradients and associated standard errors (SE) are estimated from linear regression models with fitness component (relativized at each population) against one of three traits (size PC1, forewing LD1, hindwing LD1). Estimates that are statistically significantly different from 0 at a significance level of $p < 0.05$ are shown in **bold**. N refers to sample size, Mean and SD refer to mean and standard deviation of trait (units: size PC1: $\log_e$(mm), shape LD1: centroid size) at each population.

| Sex | Locale | Trait | N | Mean | SD | β ± SE | γ ± SE |
|---|---|---|---|---|---|---|---|
| ♂ | Hoje A6 | Size PC1 | 68 | 0.24 | 0.10 | **2.06±0.99*** | -9.38±12.2 |
| | | Forewing LD1 | 68 | -2.89 | 0.67 | 0.27±0.16 | -0.054±0.29 |
| | | Hindwing LD1 | 69 | -3.09 | 0.68 | 0.11±0.16 | -0.11±0.27 |
| | Krutladan | Size PC1 | 316 | 0.22 | 0.089 | -0.38±0.52 | 11.7±7.92 |
| | | Forewing LD1 | 316 | -2.74 | 0.64 | **0.14±0.072*** | 0.047±0.14 |
| | | Hindwing LD1 | 318 | -2.95 | 0.67 | 0.044±0.069 | **-0.23±0.096*** |
| | Vombs Vattenverk | Size PC1 | 382 | 0.27 | 0.086 | -0.38±0.54 | 18.7±7.72 |
| | | Forewing LD1 | 382 | -2.68 | 0.60 | -0.029±0.077 | 0.086±0.15 |
| | | Hindwing LD1 | 384 | -2.87 | 0.65 | -0.0014±0.072 | -0.044±0.13 |
| ♀ | Hoje A6 | Size PC1 | 27 | 0.44 | 0.077 | -8.16±5.22 | -47.2±137.0 |
| | | Forewing LD1 | 27 | -2.68 | 0.53 | -1.52±0.74 | 1.80±1.60 |
| | | Hindwing LD1 | 31 | -2.75 | 0.54 | -0.079±0.79 | 0.86±1.79 |
| | Krutladan | Size PC1 | 160 | 0.39 | 0.073 | 1.55±1.24 | 47.6±29.5 |
| | | Forewing LD1 | 160 | -2.74 | 0.59 | -0.052±0.15 | 0.52±0.36 |
| | | Hindwing LD1 | 160 | -2.77 | 0.63 | -0.037±0.14 | 0.028±0.28 |
| | Vombs Vattenverk | Size PC1 | 192 | 0.39 | 0.089 | **1.83±0.68**** | **-25.7±9.80**** |
| | | Forewing LD1 | 192 | -2.58 | 0.68 | 0.075±0.090 | -0.0059±0.11 |
| | | Hindwing LD1 | 192 | -2.54 | 0.55 | 0.12±0.11 | -0.023±0.26 |

Note: * indicates $0.01 < p < 0.05$, ** indicates $0.001 < P < 0.01$, *** indicates $p < 0.001$.

Table S6. Comparison of mean-standardized linear (β) and quadratic (γ) selection gradients (estimated at population) between *I. elegans* and *E. cyathigerum*. All estimates are obtained from linear models with selection gradients (with fitness components relativized at each population) as the response variable and species identity as the predictor, weighted by inverse of standard error of selection gradients and p-values testing for the difference between two species from these models are presented.

| **Trait** | **Sex** | **β (mean estimates ± SE)** | | | **γ (mean estimates ± SE)** | | |
|---|---|---|---|---|---|---|---|
| | | *I. elegans* | *E. cyathigerum* | p | *I. elegans* | *E. cyathigerum* | p |
| Body size PC1 | ♂ | 0.26±0.73 | 0.14±0.64 | 0.87 | 0.47±10.14 | 9.27±8.99 | 0.40 |
| | ♀ | 0.74±1.57 | 0.97±1.39 | 0.89 | -8.12±13.58 | -9.40±12.19 | 0.93 |
| Wing shape LD1 (forewing) | ♂ | -0.044±0.10 | 0.099±0.089 | 0.18 | -0.003±0.14 | 0.041±0.13 | 0.78 |
| | ♀ | -0.058±0.17 | -0.082±0.15 | 0.89 | -0.049±0.20 | 0.20±0.18 | 0.24 |
| Wing shape LD1 (hindwing) | ♂ | -0.058±0.061 | 0.038±0.052 | 0.13 | -0.021±0.10 | -0.15±0.077 | 0.16 |
| | ♀ | -0.011±0.10 | 0.042±0.090 | 0.62 | -0.075±0.20 | 0.061±0.18 | 0.51 |

Table S7. Summary of models relating linear (β) selection gradients on male body size and sexual selection regime. All estimates are obtained from linear models with selection gradients as the response variable and species identity as the predictor, weighted by inverse of standard error of selection gradients. The $r^2$ of these models and p-values testing against the null hypothesis of slope = 0 are presented.

| Response | Predictor | Intercept ± SE | | Slope ± SE | p | $r^2$ |
|---|---|---|---|---|---|---|
| β (male body size PC1) | Male Density | *I. elegans* | 0.65±1.32 | -1.05±1.03 | 0.33 | <1% |
| | | *E. cyathigerum* | 1.41±1.41 | | | |
| | Female Density | *I. elegans* | 1.04±0.70 | -3.25±2.08 | 0.14 | 2.7% |
| | | *E. cyathigerum* | 0.94±0.80 | | | |
| | Operational Sex Ratio | *I. elegans* | -1.10±1.05 | **0.61±0.15** | <0.01 | 47.1% |
| | | *E. cyathigerum* | -4.90±1.33 | | | |
| | Androchrome Frequency | *I. elegans* | 2.00±0.41 | -4.43±2.57 | 0.11 | 6.0% |
| | | *E. cyathigerum* | 0.41±0.62 | | | |
| | Proportion copulating males | *I. elegans* | 1.18±0.72 | -4.50±2.37 | 0.079 | 9.2% |
| | | *E. cyathigerum* | 0.60±0.64 | | | |
| | Opportunity for sexual selection | *I. elegans* | 0.53±0.65 | **0.14±0.05** | 0.025 | 21.2% |
| | | *E. cyathigerum* | -0.70±0.77 | | | |

Table S8. Summary of models relating linear (β) selection gradients on male wing shape and sexual selection regime. All estimates are obtained from linear models with selection gradients as the response variable and species identity as the predictor, weighted by inverse of standard error of selection gradients. The $r^2$ of these models and p-values testing against the null hypothesis of slope = 0 are presented.

| Response | Predictor | | Intercept ± SE | Slope ± SE | p | $r^2$ |
|---|---|---|---|---|---|---|
| β (forewing shape LD1) | Male Density | *I. elegans* | -0.076±0.21 | 0.084±0.15 | 0.55 | 1.1% |
| | | *E. cyathigerum* | -0.073±0.17 | | | |
| | Female Density | *I. elegans* | -0.0040±0.13 | -0.20±0.35 | 0.58 | 1.2% |
| | | *E. cyathigerum* | 0.15±0.13 | | | |
| | Operational Sex Ratio | *I. elegans* | -0.22±0.24 | **0.076±0.028** | 0.016 | 34.3% |
| | | *E. cyathigerum* | -0.52±0.24 | | | |
| | Androchrome Frequency | *I. elegans* | 0.19±0.16 | -0.62±0.41 | 0.15 | 13.2% |
| | | *E. cyathigerum* | 0.14±0.089 | | | |
| | Proportion copulating males | *I. elegans* | 0.070±0.10 | -0.61±0.39 | 0.14 | 13.9% |
| | | *E. cyathigerum* | 0.16±0.094 | | | |
| | Opportunity for sexual selection | *I. elegans* | -0.17±0.097 | 0.017±0.09 | 0.077 | 19.9% |
| | | *E. cyathigerum* | -0.065±0.12 | | | |
| | Forewing shape LD1 | *I. elegans* | 1.07±0.96 | **-0.51±0.19** | 0.019 | 32.5% |
| | | *E. cyathigerum* | -1.30±0.54 | | | |
| β (hindwing shape LD1) | Male Density | *I. elegans* | -0.11±0.099 | -0.021±0.091 | 0.82 | 2.4% |
| | | *E. cyathigerum* | 0.064±0.124 | | | |
| | Female Density | *I. elegans* | 0.015±0.058 | -0.31±0.20 | 0.15 | 16.1% |
| | | *E. cyathigerum* | 0.11±0.071 | | | |
| | Operational Sex Ratio | *I. elegans* | -0.13±0.22 | 0.031±0.018 | 0.11 | 18.9% |
| | | *E. cyathigerum* | -0.22±0.16 | | | |
| | Androchrome Frequency | *I. elegans* | -0.062±0.10 | 0.0098±0.26 | 0.97 | 2.1% |
| | | *E. cyathigerum* | 0.037±0.057 | | | |
| | Proportion copulating males | *I. elegans* | -0.017±0.065 | -0.22±0.25 | 0.39 | 7.2% |
| | | *E. cyathigerum* | 0.061±0.059 | | | |
| | Opportunity for sexual selection | *I. elegans* | -0.129±0.058 | 0.0098±0.0052 | 0.082 | 21.7% |
| | | *E. cyathigerum* | -0.057±0.070 | | | |
| | Hindwing shape LD1 | *I. elegans* | 0.27±0.87 | -0.14±0.16 | 0.41 | 7.0% |
| | | *E. cyathigerum* | -0.38±0.49 | | | |

Table S9: Estimates for quantitative parameters of social organization in *I. elegans* and *E. cyathigerum* obtained from mixed effect models with one of six parameters as the response variable, species identity as the fixed effect and sampling year as the random effect. Mean estimates for each species calculated from outputs of mixed effect models and p-values testing against the null hypothesis of no difference between species are presented. (Model: one of 6 parameters describing social organization ~ Species + (1|Locale) + (1|Year))

| **Parameter** | **Estimate (Mean ± SE)** | | **p-value** |
|---|---|---|---|
| | *Ischnura elegans* | *Enallagma cyathigerum* | |
| Male Density | 0.316 ± 0.044 | 0.328 ± 0.046 | 0.788 |
| Female Density | 0.188 ± 0.020 | 0.081 ± 0.021 | < 0.001 |
| Operational Sex Ratio | 2.868 ± 0.846 | 8.721 ± 0.746 | < 0.001 |
| Androchrome Frequency | 0.457 ± 0.048 | 0.280 ± 0.115 | < 0.001 |
| Proportion copulating males | 0.133 ± 0.023 | 0.091 ± 0.018 | 0.067 |
| Opportunity for sexual selection | 9.759 ± 2.309 | 14.478 ± 2.126 | 0.044 |

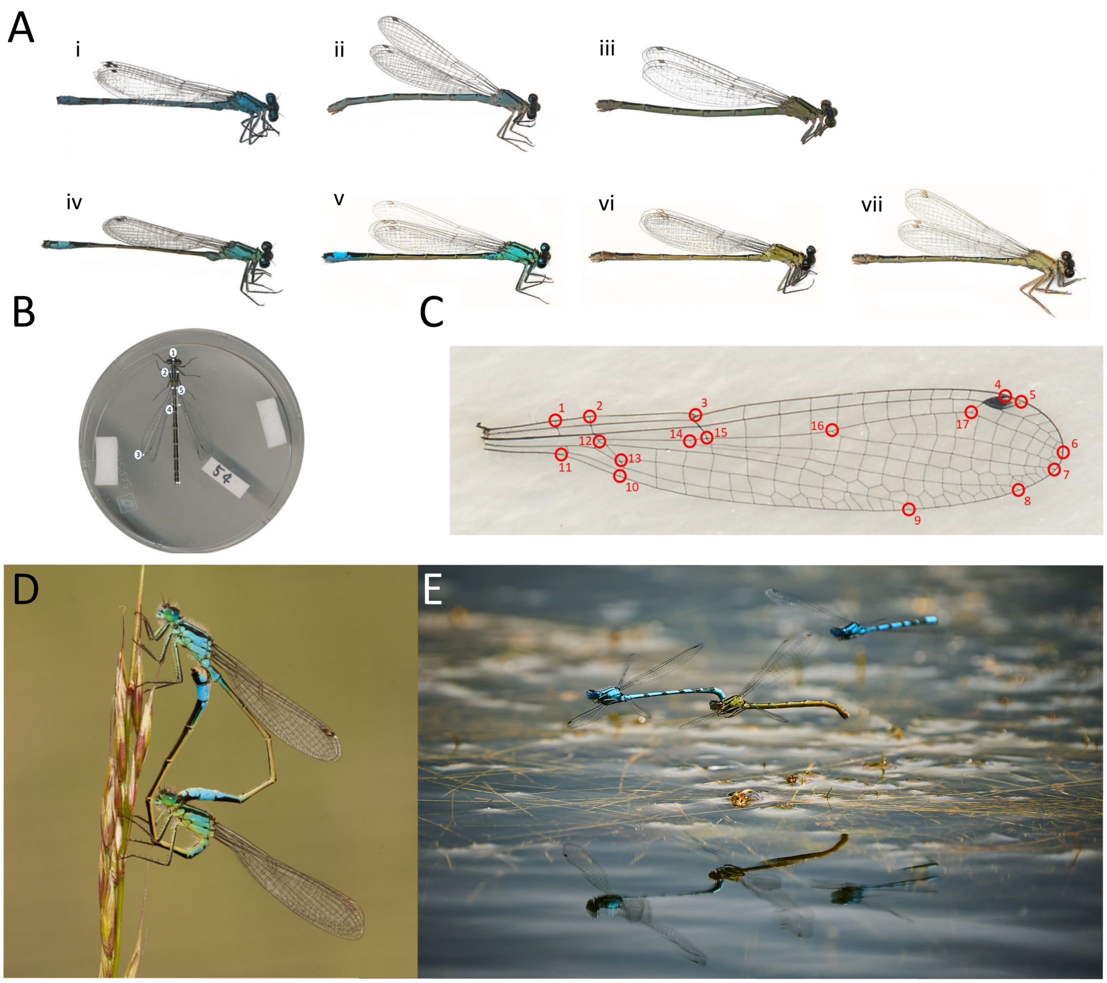


Figure S1: A) Males and female phenotypes and different color morphs in *E. cyathigerum* (i to iii) and *I. elegans* (iv to vii). Each subpanel shows an image of the following phenotype: males (i, iv), androchrome female (ii, v), gynochrome female (iii), infuscans female (vi), *infuscans-obsoleta* female (vii). Note that in both *E. cyathigerum* and *I. elegans* there is a male-mimicking female morph (androchrome females) but in former species there is only one gynochrome color morph, whereas in the latter there are two (*infuscans* and *infuscans-obsoleta*, respectively). B) Morphometric measurements of a female *E. cyathigerum* (all other males and females of *E. cyathigerum* and *I. elegans* were also measured similarly); 1: body length, 2: thorax width, 3: wing length, 4: width of abdomen segment S4, and 5: total abdomen length; C) 17 landmark position on a left forewing of a male *E. cyathigerum* that were used for estimation of the wing shape (all other males and females of *E. cyathigerum* and *I. elegans* were also landmarked similarly). D) a mating pair of *I. elegans* forming a mating wheel. E) a pair of *E. cyathigerum* forming a tandem.

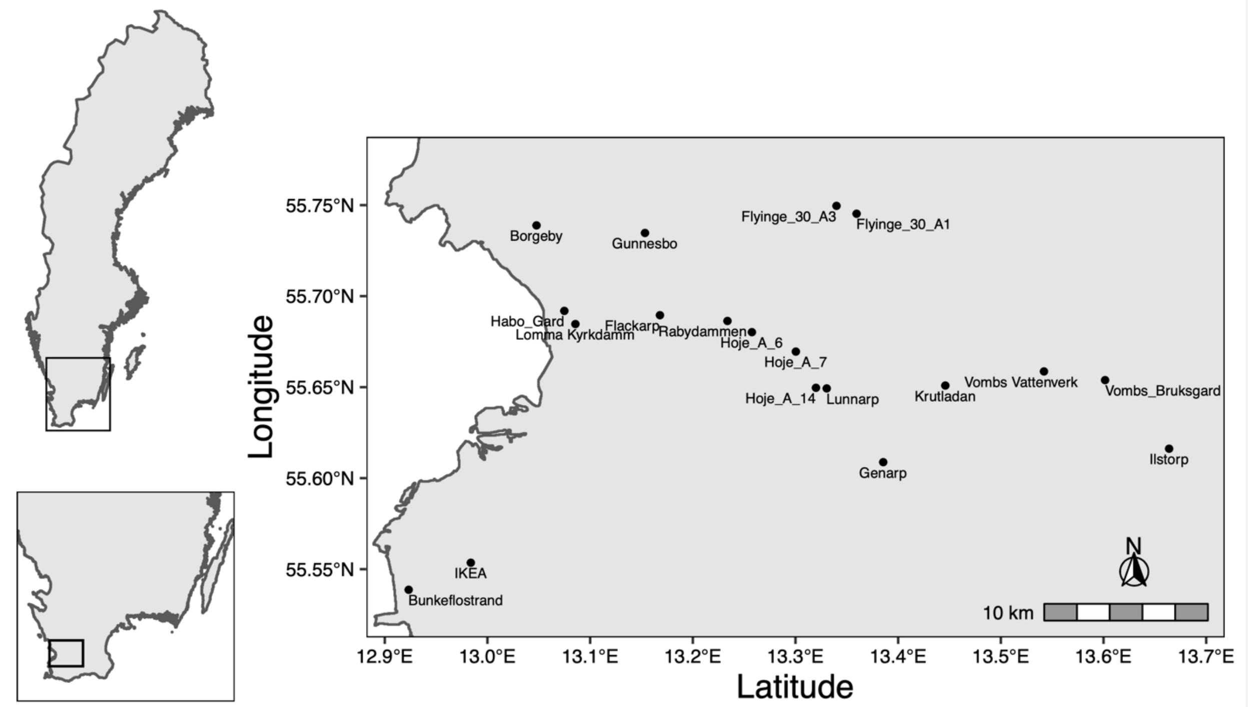


Figure S2: Geographic location of field sites in Skåne in Sweden where sampling of *E. cyathigerum* and *I. elegans* was carried out.

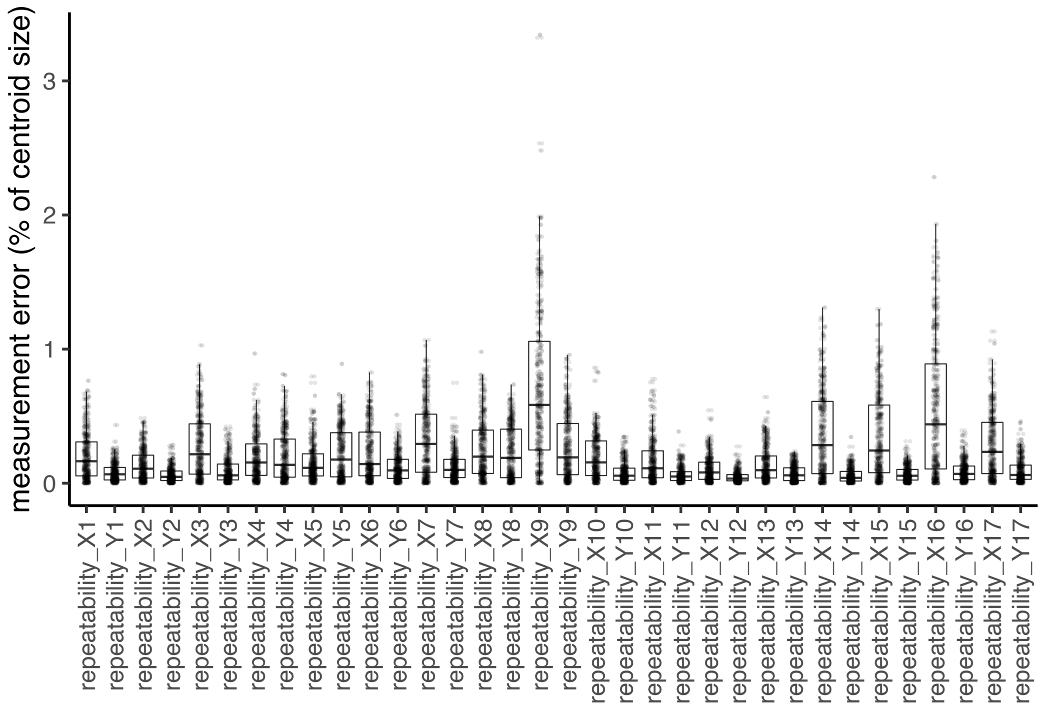


Figure S3. Repeatability of aligned shape coordinates with respect to difference in images. Difference between aligned coordinates obtained from two independently obtained images are presented as % of centroid size and compared across traits. Average measurement errors±SE: 0.194±0.002, range: 0 - 3.343%.

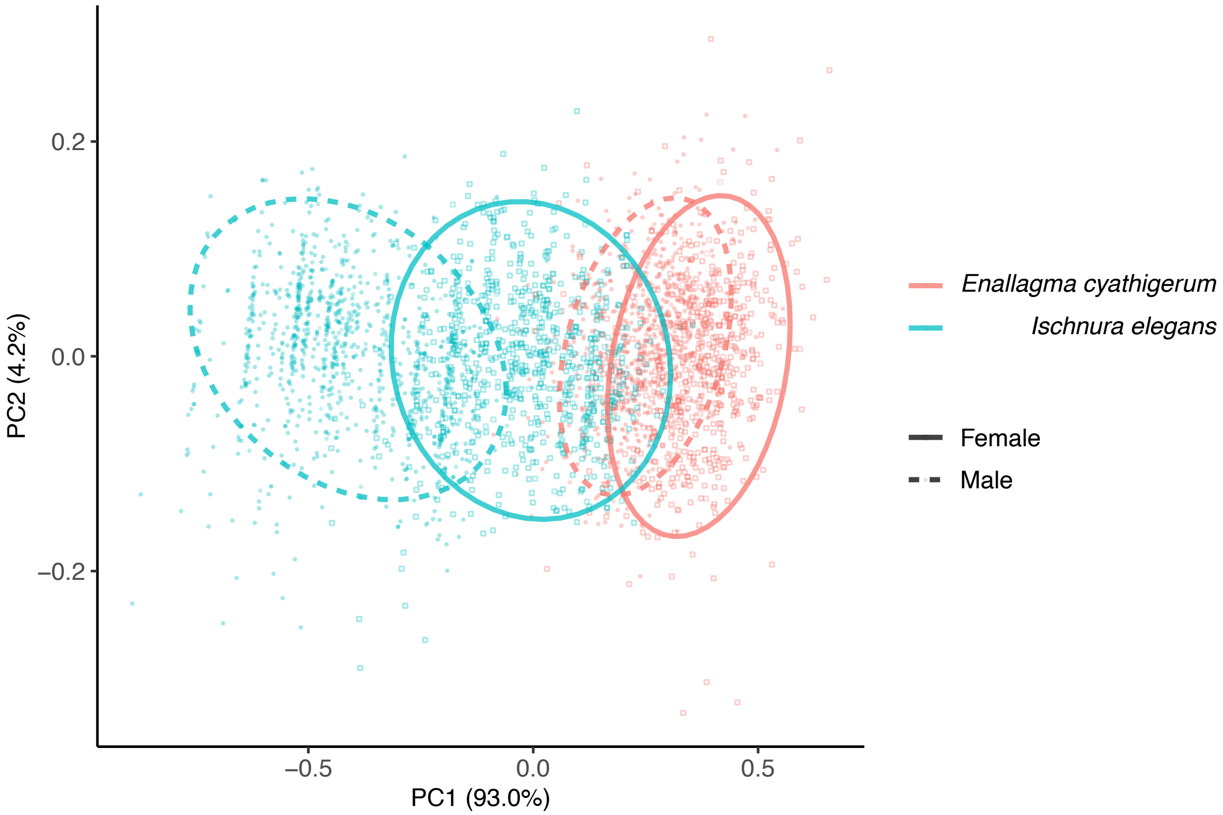


Figure S4: Principal component analysis (PCA) run on the covariance matrix of natural logarithmic values of the five size traits (body length, thorax width, abdomen length, wing length, and width of the abdominal segment S4). Ellipsoids represent 95% confidence limits for each species and each sex.

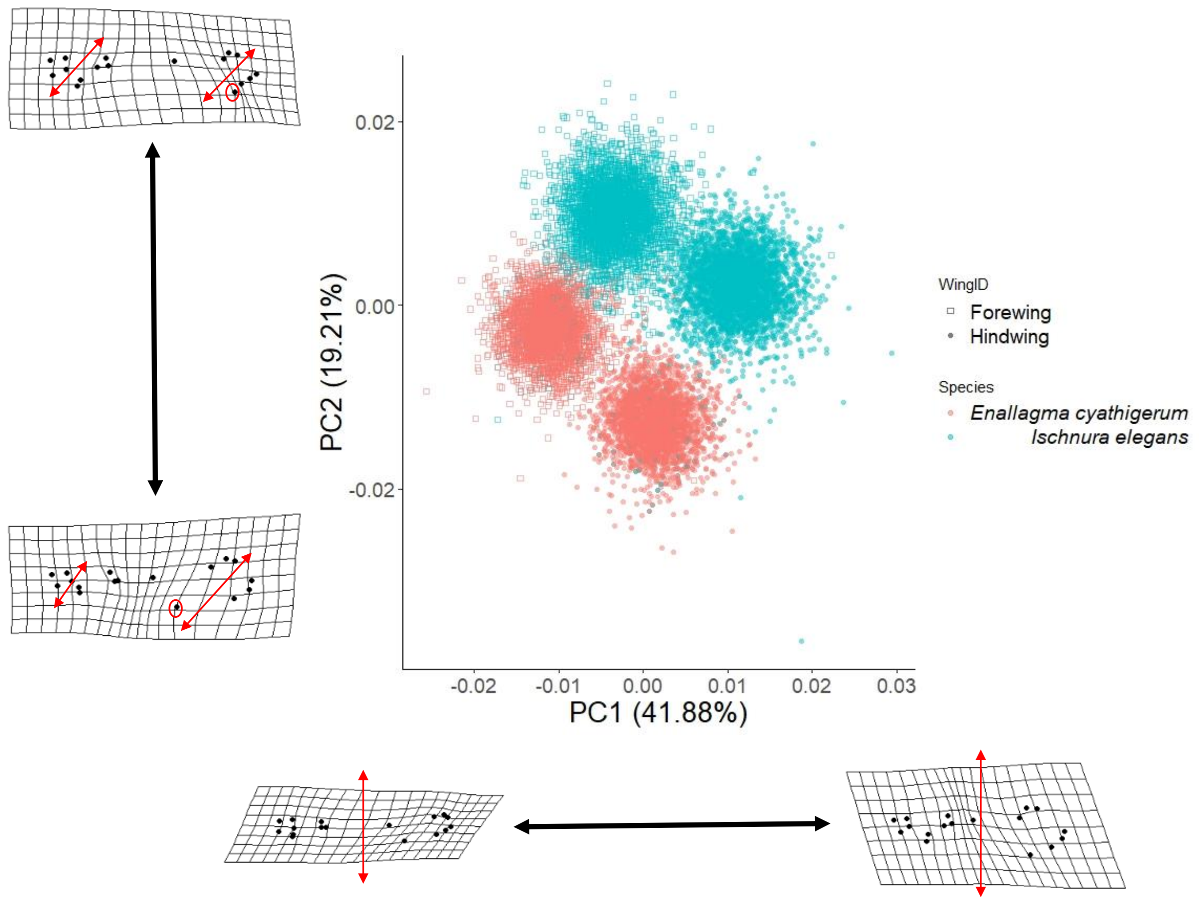


Figure S5: Wing shape variation along Principal Components, PC1 and PC2 across *E. cyathigerum* and *I. elegans*. The landmark specimens visualized in this figure were made by exaggerating the minimum and maximum PC values by a factor of two.

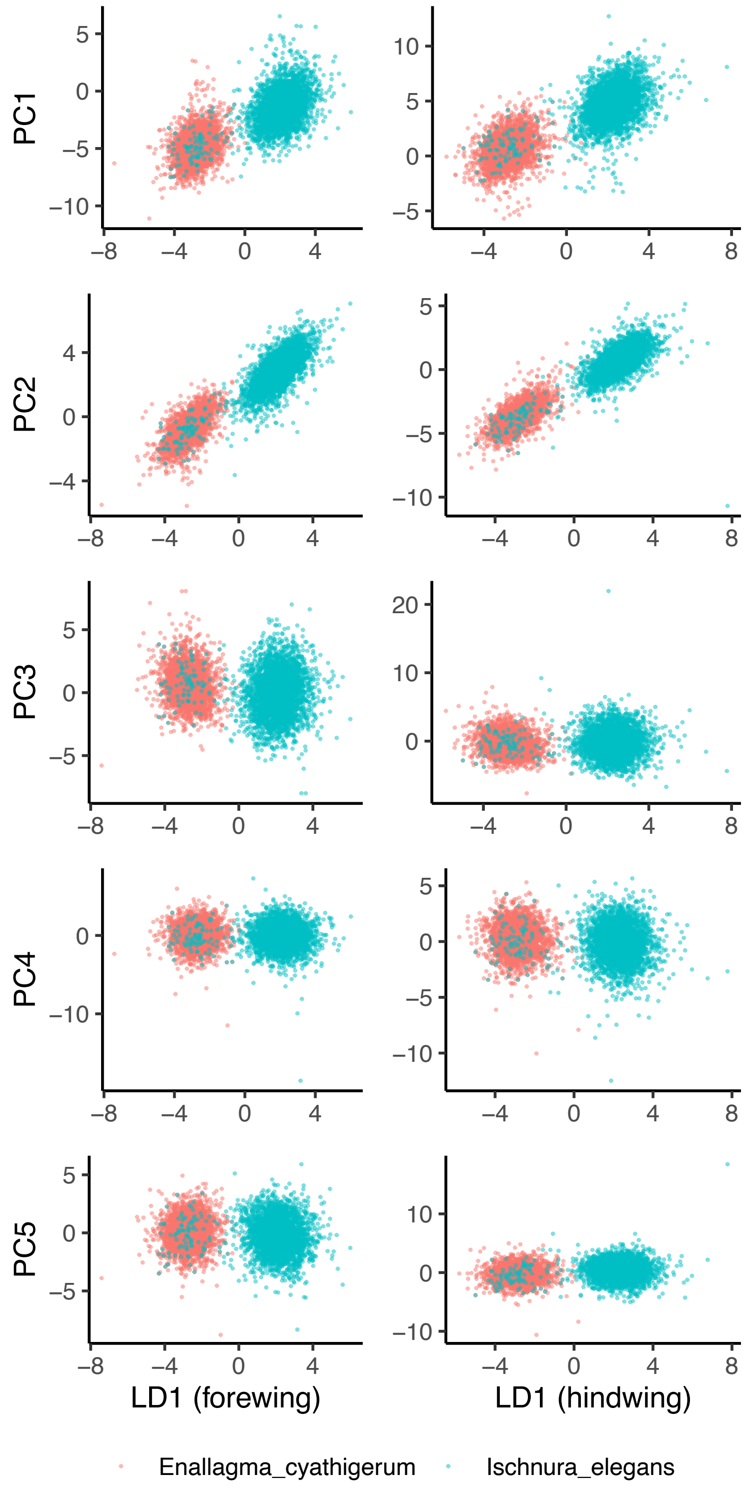


Figure S6. Relationship between principal components and linear-discriminant axis describing species differences (LD1). Left column represents forewings and right columns represents hindwings. The $r^2$ of presented relationships are as follows. PC1: forewing: 62.5%, hindwing: 68.6%, PC2: forewing: 86.9%, hindwing: 89.0%, PC3: forewing: 3.9%, hindwing: < 1%, PC4: forewing: < 1%, hindwing: < 1%, PC5: forewing: < 1%, hindwing: 4.1%.

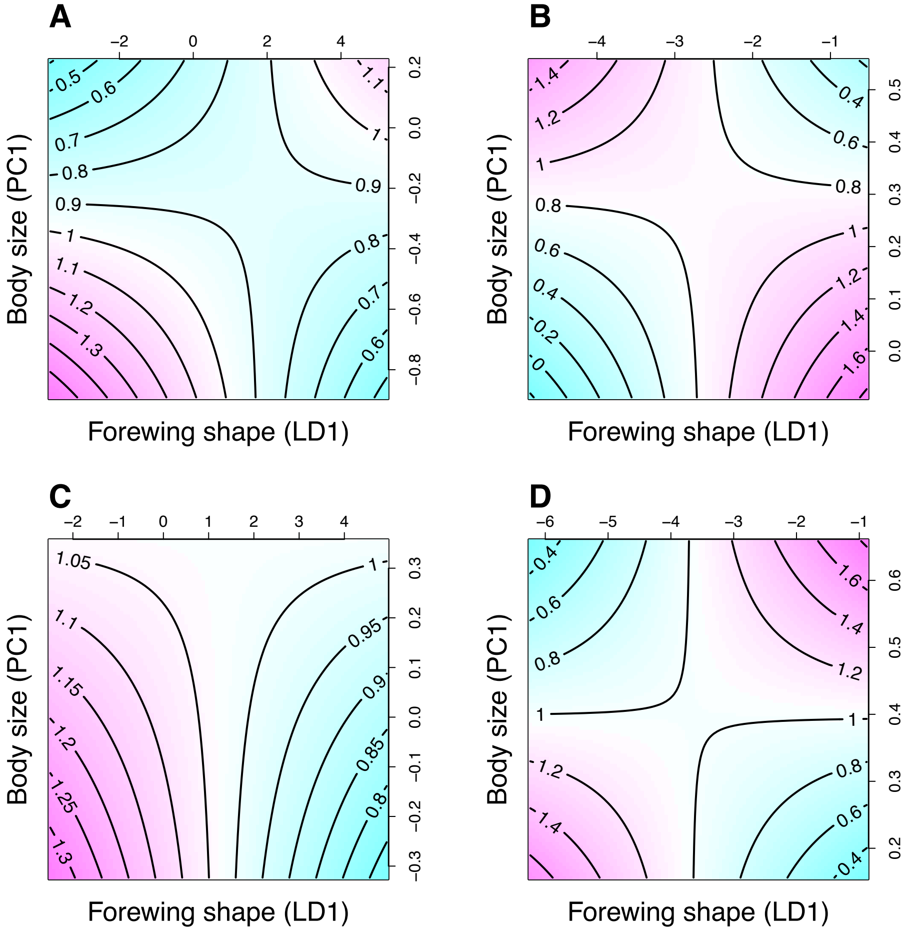


Figure S7. Bivariate fitness surfaces of forewing shape (LD1) and body size (PC1) in A) males of *I. elegans*, B) males of *E. cyathigerum*, C) females of *I. elegans* and D) females of *E. cyathigerum.* The axis perpendicular to the presented plane represents relative fitness, which is the predicted probability of being found as mated in male (A and B) and the predicted number of eggs laid by females (C and D). These fitness surfaces are based on splines from multivariate generalized additive models. The selection coefficients from this model are presented in Table 2.

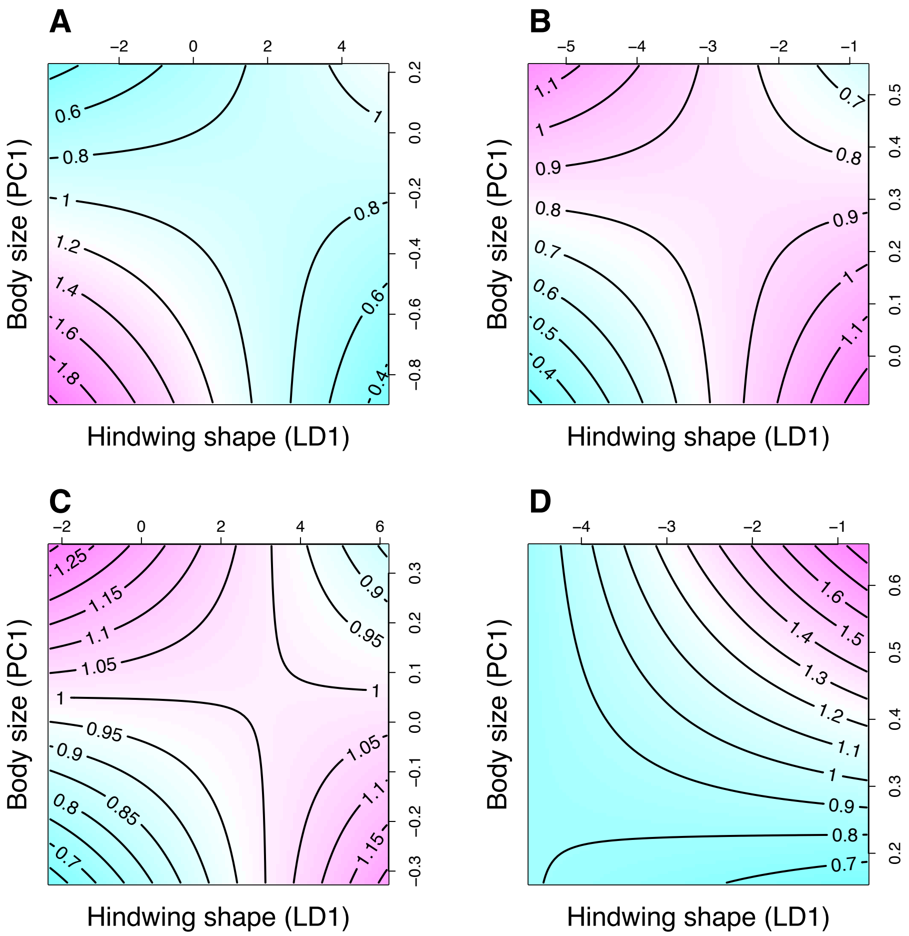


Figure S8: Bivariate selection surface of hindwing shape (LD1) and body size (PC1) in A) males of *I. elegans*, B) males of *E. cyathigerum*, C) females of *I. elegans*, D) females of *E. cyathigerum.* The axis perpendicular to the presented plane represents relative fitness, which is the predicted probability of being found as mated in male (A and B) and the predicted number of effs laid in female (C and D) based on splines evaluated with multivariate generalized additive models. The coefficients of selection in this model are presented in Table 3 and the statistical analysis for their calculation is presented in Table S1 and S2 in the supplementary information.

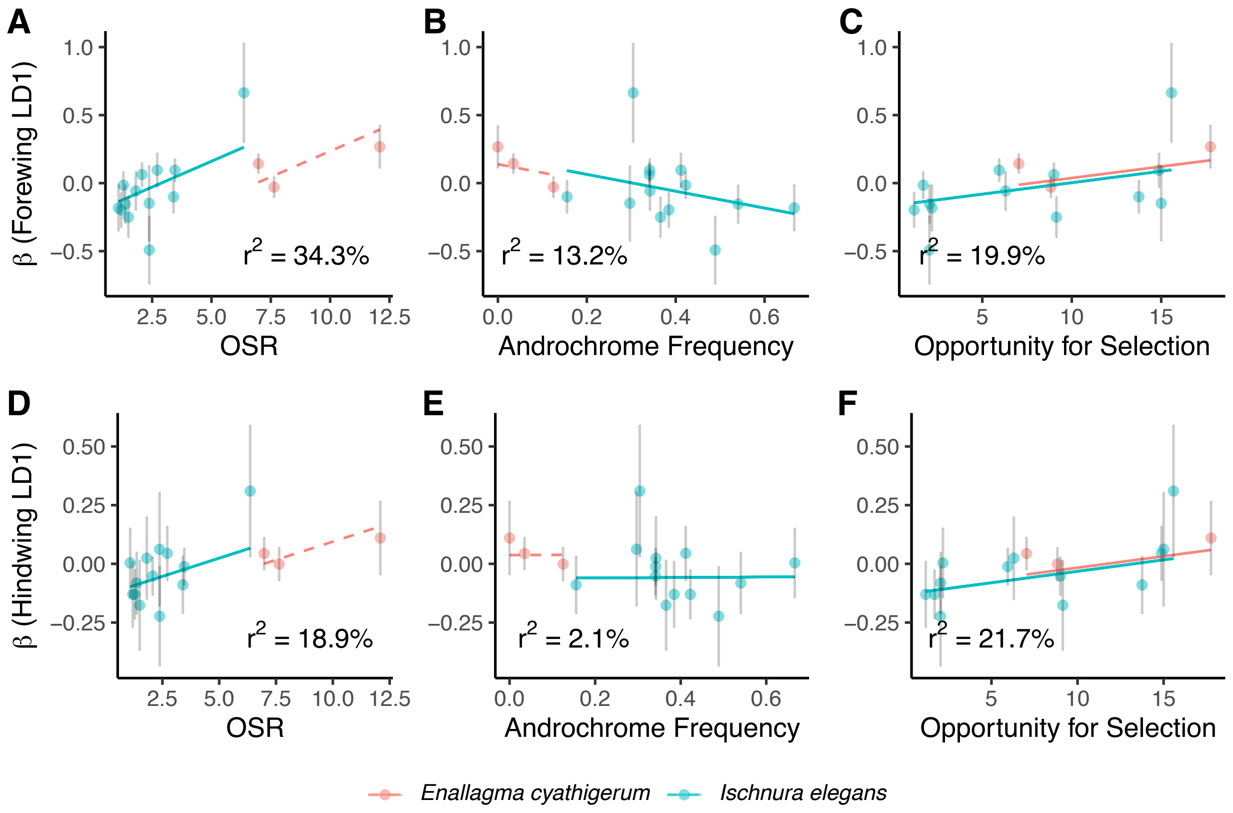


Figure S9. Relationship between directional selection gradient on male forewing shape (LD1) and (A) operational sex ratio (OSR), (B) androchrome frequency, and (C) opportunity for selection, between directional selection gradient on male hindwing shape (LD1) and (D) OSR, (E) androchrome frequency, and (F) opportunity for selection. Each point represents a population with unique species-locale-year identity and error bar shows standard errors of selection gradients. Regression lines are derived from a weighted regression model (shown in Table S8). Note that one population (*I. elegans* from Hoje A7 in 2021, see Table S3 and S4 for estimates of this population) was removed from plots for better visualization, while the regressions are from analyses using the whole dataset.

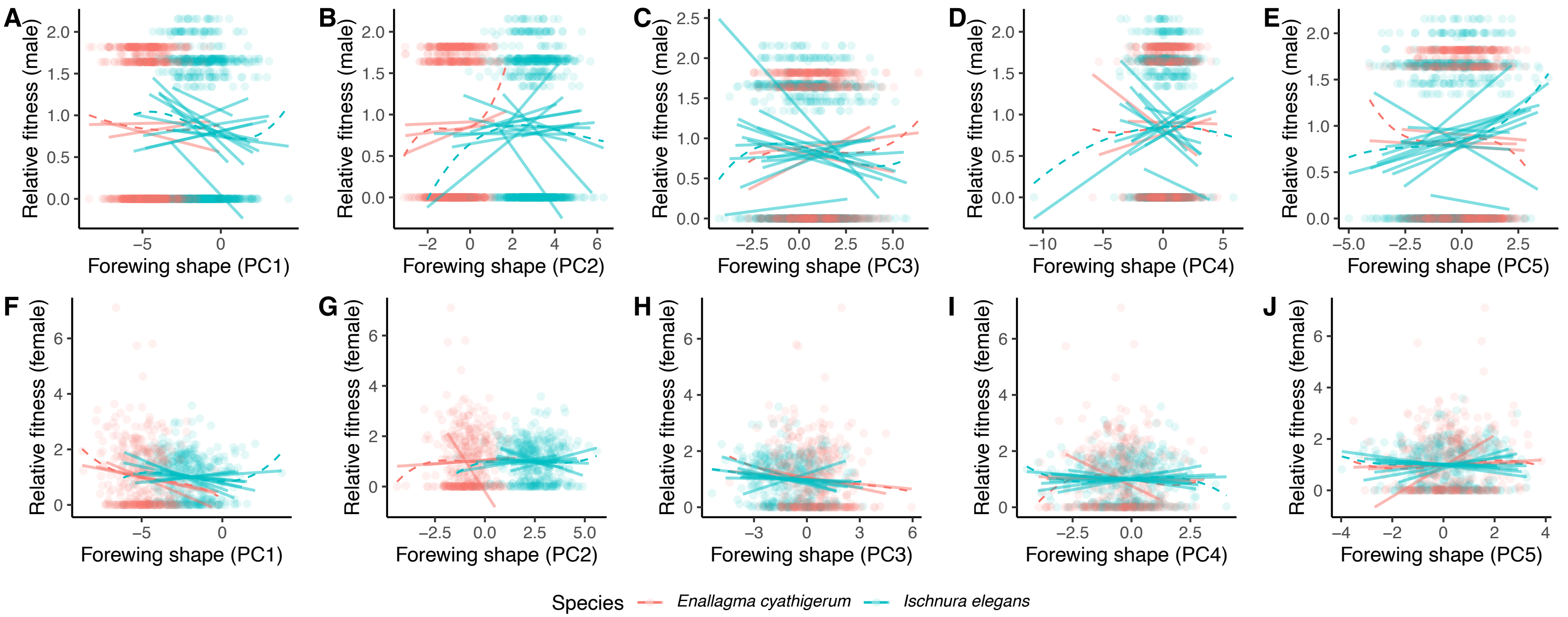


Figure S10: Comparison of species-specific fitness surface with directional selection gradients at each locality (3 populations in *E. cyathigerum* and 14 populations in *I. elegans*). Sexual selection (males, A-E) and fecundity selection (females, F-J) on fore wing shape (PC1-PC5) are plotted. Fitness functions of each species (dashed lines) were visualized using cubic splines. Fitness data on the Y-axis were relativized at each location. Directional selection gradients (solid lines) were estimated using a regression model of relative fitness against shape at each location.

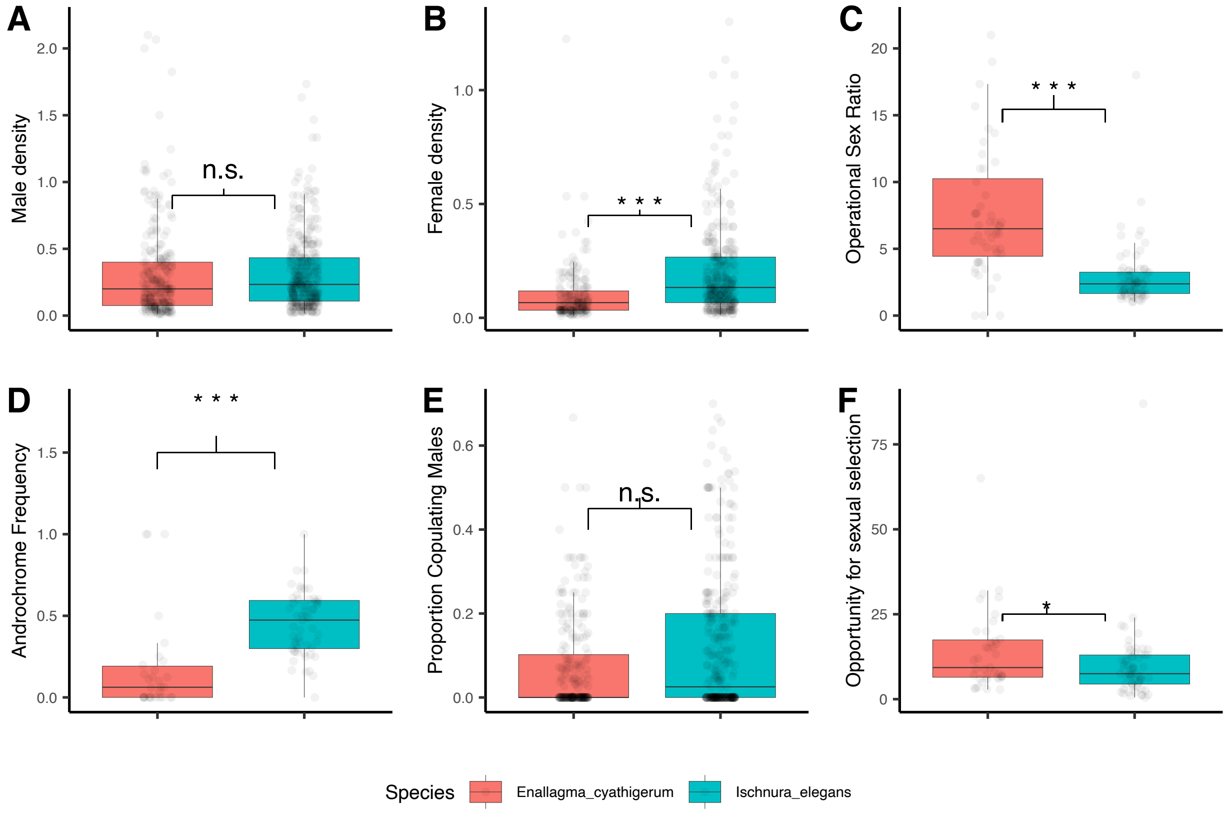


Figure S11: Boxplots (barline in the boxplot represent 50th percentile and the intervals represent the range between 1.5 times above 75th percentile and below 25th percentile) comparing demographic and mating system parameters of *E. cyathigerum* and *I. elegans* across four consecutive years (2018, 2019, 2020, 2021) of sampling. A) Male density, B) female density, C) operational sex ratio, D) androchrome frequency, E) proportion of copulating males, and F) opportunity for sexual selection. The estimates for the mean and standard errors of these parameters are as well as the statistical analysis for comparison between the two species is presented in Table S9 in the supplementary information. Symbols of significance: ***$P < 0.001$, **$P < 0.01$, *$P<0.05$.
